# The biofilm matrix scaffold of *Pseudomonas* species contains non-canonically base paired extracellular DNA and RNA

**DOI:** 10.1101/527267

**Authors:** Thomas Seviour, Fernaldo Richtia Winnerdy, Lan Li Wong, Xiangyan Shi, Sudarsan Mugunthan, Remi Castaing, Sunil S Adav, Gurjeet Singh Kohli, Heather M Shewan, Jason R Stokes, Scott A. Rice, Anh Tuân Phan, Staffan Kjelleberg

## Abstract

While extracellular DNA (eDNA) is recognized as a critical biofilm matrix component, it is not understood how it contributes to biofilm function. Here we isolate eDNA from *Pseudomonas* biofilms using ionic liquids, and discover that its key biophysical signatures, i.e. fluid viscoelasticity, nucleic acid conformation, and temperature and pH dependencies of gel to solution transitions, are maintained. Solid-state analysis of isolated eDNA, as a proxy for eDNA structure in biofilms, revealed non-canonical Hoogsteen base pairs, triads or tetrads involving guanine and thymine or uracil. These were less abundant in chromosomal DNA and undetected as eDNA underwent gel-sol transition. Purine-rich RNA was present in the eDNA network, which potentially enables eDNA to be the main cross-linking exopolymer in the matrix through non-canonical nucleobase interactions. Our study suggests that *Pseudomonas* assemble extracellular DNA and RNA into a network with viscoelastic properties, which underpin their persistence and spreading, and may aid the development of more effective controls for biofilm-associated infections.

## Introduction

Biofilms are key microbial ecosystems that contribute to bacterial pathogenicity (1), disrupt flow in water filtration systems (2) and facilitate wastewater treatment bioprocesses (3). They represent bacterial adaptation strategies allowing for increased antibiotic tolerance (4), enhanced resource capture (5) and the establishment of ecological microniches (6). Such properties are unique to biofilms, in contrast to planktonic bacteria, and are not mediated directly by the cells but instead by an extracellular polymeric matrix the cells secrete (7). While cells themselves are stiff and rigid, the matrix provides the biofilm with a viscoelastic property that allows cells to withstand mechanical and chemical stress, and enhances surface colonization by facilitating biofilm migration. Moreover, the viscoelastic property of the matrix itself has been identified as a virulence factor in chronic biofilm infections (8).

Exopolymer functions in biofilms have been studied extensively, particularly for *Pseudomonas aeruginosa*, which contributes to one in five clinical infections (9). No fewer than eight exopolymers have been identified as supporting key traits in *P. aeruginosa* biofilms, including three exopolysaccharides (10), four proteins (11–13) and extracellular DNA (eDNA) (14). While progress has been made towards describing the structures and identities of extracellular polysaccharides and proteins, by applying classical chemical and molecular approaches, the important question regarding extracellular DNA is not what it is but how it differs structurally from chromosomal DNA and what enables it to perform a structural function in the biofilm matrix. eDNA has been described as a key matrix biopolymer in clinical (15) and environmental biofilms (16), particularly in *P. aeruginosa* biofilms (13), and has been shown to colocalize with putative *P. aeruginosa* polysaccharide Pel (17) and be degraded by DNA-specific endonucleases (18). While there have been attempts to explain the properties of eDNA, they have tended to focus on primary-structure differences with the chromosomal DNA and as yet, no signature eDNA sequences have been identified (19).

DNA can form multiple higher-order structures arising from differences in torsional stress as the two strands are twisted around the axis. For chromosomal DNA (cDNA), this is known to consist of supercoiled duplex structures that are formed by the action of histone-like proteins and topoisomerases, and then relaxed by gyrases to allow replication and transcription to occur (20). DNA supercoiling, however, does not reconcile with descriptions of eDNA as a primary biofilm structural agent which would imply a networked structure. While eDNA has been shown to colocalize with DNA-binding proteins (21), this does not distinguish it from chromosomal DNA nor allude to its higher-order structure. We propose that a molecular understanding of how the eDNA is assembled and organized is key to answering how and why DNA transforms from the chromosomal form to that found in the biofilm matrix. In this study, we elucidate molecular interactions that lead to the higher-order structure of eDNA. We seek to contrast the biophysical properties of eDNA and chromosomal DNA and correlate differences to a distinct eDNA molecular signature. We hypothesize that isolating eDNA such that its higher-order structure is preserved should enable us to describe interactions in eDNA that underpin its ability to form networked structures.

## Results

### eDNA elasticity is preserved during extraction

To isolate eDNA from *P. aeruginosa* biofilms while preserving its higher-order structure, we use ionic liquid 1-ethyl-3-methyl-imidazolium acetate (EMIM-Ac). This has been previously shown to non-destructively dissolve a range of recalcitrant biopolymers, including DNA (22) and cellulose (23), and we recently demonstrated its use in dissolving *P. aeruginosa* biofilm exopolymers (24). Here, we consider the change in fluid rheology upon dissolving *P. aeruginosa* biofilms in EMIM-Ac, whereby we observe the subsequent fluid to be highly viscoelastic which is characteristic of high molecular weight polymers in solution. We observe viscoelasticity, in that the EMIM-Ac following biofilm dissolution displayed both elastic and viscous characteristics, upon measuring a normal force during viscosity measurements on a parallel plate rheometer as a function of shear rate under steady shear conditions. The viscosity is only slightly shear-thinning (Supplementary Figure 1A and Supplementary Table 1). Normal force (i.e. contact force perpendicular to the parallel plates applying shear), however, is zero for Newtonian fluids and thus the observation of finite values indicates the presence of an anisotropic polymer in the fluid, giving rise to non-linear elasticity, which is expected for dilute polymer solutions in viscous Newtonian fluids (i.e. Boger fluids) (25). Normal force is characterized as a normal stress difference (*ΔN* = *N*_1_-*N*_2_), where *N*_1_ and *N*_2_ are primary and secondary normal stress differences respectively (see definition in Materials and Methods (26)). The fluid’s elasticity is observed to dominate over the viscous flow properties for the wild type biofilm in EMIM-Ac, whereby *ΔN* is an order of magnitude greater than shear stress (Figure 1A; Wild type). The solvent (EMIM-Ac) alone exhibited no elasticity, indicating that the elastic properties arise from the dissolution of polymeric components within the biofilm matrix. *ΔN* has a power law dependence on the shear rate with an exponent (*n*) of 1.4 (Supplementary Figure 1B, Wild type). We find that the viscosity and non-linear elasticity measurements are accurately modeled using the modified, finitely-extensible nonlinear elastic (FENE-P) polymer model (Figure 1A-B; Supplementary Tables 1 and 2), which is commonly used to describe solutions containing flexible/semi-flexible polymers using the common assumption that *ΔN* ~ *N*_1_ (27). As DNA is a very high molecular weight semi-flexible polymer, we hypothesize that this may be the dominant polymer giving rise to the nonlinear elastic properties of the fluids.

**Table 1:** Relative abundances of monoribonucleotides in extracellular eDNA gel isolate from *Pseudomonas aeruginosa* biofilm as determined by integrating 31P NMR spectrum following alkalinization.

| $^{31}\text{P}$ shift (ppm) | Ribonucleotide | Relative abundance |
| --- | --- | --- |
| 4.03 | 3' AMP | 18 % |
| 4.01 | 3' GMP | 17 % |
| 3.90 | 3' CMP/3' UMP | 9 % |
| 3.87 | 3' CMP/3' UMP | 11 % |
| 3.68 | 2' CMP/2' UMP | 10 % |
| 3.66 | 2' CMP/2' UMP | 12 % |
| 3.50 | 2' AMP | 10 % |
| 3.42 | 2' GMP | 13 % |

**Figure 1:**
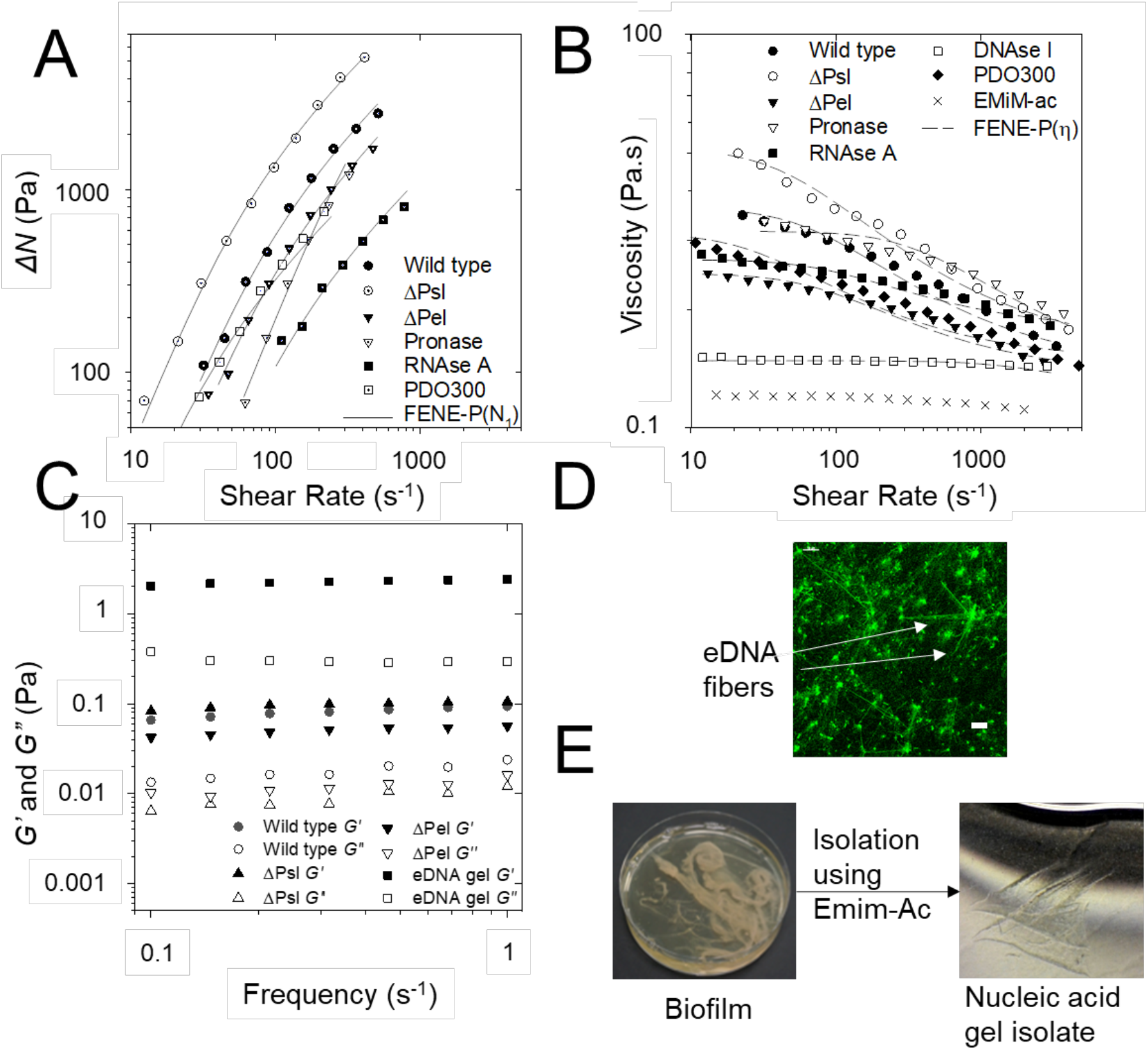
(A) Non-linear elasticity as normal stress difference (*ΔN* = **N*_1_-*N*_2_*) and (B) viscosity against shear rate for *Pseudomonas aeruginosa* biofilm wild type, PDO300, Δ*psl*, Δ*pel*, and pronase, RNase A and DNase I digested wild type biofilm in 1-ethyl-3-methylimidazolium acetate (40 mg/mL) at 25 °C, 250 μm gap. This is measured as a function of shear stress from 10 to 1000 Pa. *ΔN* is not described for DNase I digested biofilm in Figure 1A and Supplementary Figure 1B as their normal force (F_N_) is less than the resolution of the rheometer (i.e. 0.1 N) and set to zero for calculating *ΔN*. Both the *ΔN* and viscosity data are fitted with the FENE-P model, a rigid dumbell model for polymer solutions. Fitting parameters are shown in Supplementary Table 2. (C) Frequency dependence rheogram showing that storage (*G*′ or elastic) exceeds loss (*G*′′ or viscous) modulus for wild type, Δ*psl* and Δ*pel* biofilms and isolated eDNA gel at 25°C (250 μm plate gap). (D) Micrograph of *P. aeruginosa* biofilm DNA stained green with TOTO-1 (scale bar 10 μm). (E) Phase separation of extracellular DNA extracted from *P. aeruginosa* biofilms into a gel occurs upon transfer from 1-ethyl-3-methylimidazolium acetate (EMIM-Ac) into water.

The polysaccharides alginate, Pel and Psl have been suggested as the structural scaffolds of *P. aeruginosa* biofilms (10). To investigate the possibility that the polysaccharides, rather than the DNA, cause the elastic behavior of the fluid after biofilm dissolution, an alginate over-expressing strain PDO300 (i.e. mucoid) and its isogenic Pel and Psl genetic knockout mutants were dissolved in EMIM-Ac (28). When Psl is absent, the elasticity (i.e. *ΔN*) and viscosity increase relative to the wild type, and the mucoid strain PDO300 is less elastic than the wild type despite the over-expression of alginate (Figure 1; Δ*psl*). In contrast, when Pel is absent, there was a slight decrease in both elasticity and viscosity relative to both the wild type and PDO300 (Figure 1; Δ*pel*).

The contributions of proteins, RNA and DNA were also explored by digestion using pronase, RNase A and DNase I, respectively (Figure 1; Pronase, RNase, DNase). Removing each exopolymer component individually decreased biofilm elasticity in EMIM-Ac. All treatments, except for DNase I-treated biofilm, displayed *n* values of 0.9 – 1.7 (Supplementary Table 1), characteristic of elasticity imparted by DNA (29). While viscosity was unchanged following pronase treatment, there was a slight decrease in elasticity. This decrease in elasticity was even greater following RNase treatment and elasticity was completely lost upon DNase treatment, with *ΔN* reduced to zero. We suggest that the elastic response of the biofilm in EMIM-Ac can thus be primarily attributed to DNA. Elasticity was not detected in EMIM-Ac solutions of calf thymus DNA at the same concentration, indicating that elasticity is not a universal property of DNA solutions in EMIM-Ac.

Furthermore, characterization of linear viscoelastic properties of hydrated native *P. aeruginosa* biofilms (i.e. in water rather than EMIM-Ac) under conditions of oscillating shear show that the storage (or elastic) modulus (*G*′) is greater than the loss (or viscous) modulus (*G*′′) across the frequency range 0.1 to 1 s^−1^, even in the absence of Pel and Psl (Figure 1C), providing further evidence that they are not primarily responsible for the viscoelastic property of the biofilm. This is consistent with the hydrated biofilm being classified as a gel. A storage modulus could not be measured following digestion of the native biofilm by DNase I, and biofilm digested with heat-inactivated DNase I also remained as a gel, illustrating that the lack of measurable G’ was due to enzymatic activity (Supplementary Figure 1C).

DNA is thus considered the main contributor to the elastic gel-like rheology of *P. aeruginosa* biofilms, and to the non-linear elasticity upon dissolution of lyophilized *P. aeruginosa* biofilm in EMIM-Ac. The fundamental structure of the DNA within the two systems is different; a gel-like rheology may arise from a polymer network structure, while the non-linear elasticity indicates the movement and deformation of high molecular weight polymers during flow. The approach of dissolving biofilm in EMIM-Ac therefore allows the assessment of the contribution of individual exopolymers to its properties that is complemented by rheological assays on the native biofilms.

The data suggest that multiple exopolymers contribute to the rheology, with the matrix from the Δ*psl* mutant being the most elastic, followed by wild type and RNase A-treated biofilms, which were the least elastic. It has been noted previously that a range of *P. aeruginosa* exopolymers influence biofilm extracellular matrix crosslinking (10) and the data presented here further clarify that these exopolymers can only modulate biofilm rheology if eDNA is present. An eDNA scaffold is therefore a prerequisite for this rheological differentiation of the matrix. Further support for the presence of an eDNA scaffold is provided in the micrograph of a biofilm stained with TOTO-1 for DNA visualization (Figure 1D), which shows DNA fibers in the extracellular matrix.

The absence of phospholipids and lipopolysaccharides in EMIM-Ac following biofilm dissolution demonstrate that EMIM-Ac did not lyse either biofilm or planktonic cells (Supplementary Figure 2A). Therefore, it was concluded that the DNA dissolved following treatment of *P. aeruginosa* biofilms with EMIM-Ac is extracellular and not due to extraction of intracellular DNA.

### Key biophysical signatures of nucleic acids are preserved during isolation

We recovered the eDNA from EMIM-Ac following biofilm dissolution by exploiting the ability of perchloric acid to selectively precipitate DNA over protein (Supplementary Figure 2B). Following further purification by gel permeation chromatography, the polymer phase-separated into a gel upon transfer from EMIM-Ac into water (i.e. the gel isolate), mimicking the formation of networks in native biofilms (Figure 1E). *G*′ exceeded *G*′′ across the same frequency range as for the hydrated, native *P. aeruginosa* biofilms, consistent with gels, as was the case for the biofilms (Figure 1C) and as distinct from the viscous behavior (G’ < G’’) commonly displayed by aqueous solutions of DNA (30). Furthermore, DNase degraded the isolated gel into shorter DNA fragments (Supplementary Figure 3A). Other fractions, including those not precipitated by perchloric acid, did not self-assemble into gels (Supplementary Figure 3B). Similarly, calf thymus DNA did not form gels when processed the same way, either with or without added cations (Supplementary Figure 3C) suggesting, once again, that networking and elastic behavior is not a general feature of DNA under the conditions applied in this study. Moreover, the biofilm and isolated eDNA displayed similar phase transition temperatures of 56.8 and 57.2°C, respectively (Supplementary Figures 4A and 4B), and the elasticity described for the isolated eDNA and biofilm in Figure 1C was lost at temperatures exceeding the phase transition temperatures and upon alkalinization (i.e. 0.2 M NaOH).

Circular dichroism (CD) is a highly sensitive spectroscopic technique for determining the secondary structure of biomolecules, particularly proteins and nucleic acids. It was used here to understand whether the nucleic acid (NA) conformation was modified during extraction and isolation. Unprocessed *P. aeruginosa* biofilms displayed a major circular dichroism (CD) peak at 250-285 nm (Figure 2A), which is consistent with the presence of NA (31) and this peak was also observed to dominate the CD spectrum of the gel isolate (Figure 2B). The spectral trough at 225.5-226.5 nm is typical for proteins (32). NA can also display a trough in this region, although the relative depth of the trough and its absence after proteolysis and fractional precipitation that excludes proteins (Supplementary Figure 2B) suggest that it denotes proteinaceous material. Nonetheless, while the CD spectrum shows that proteins are significant components of the biofilm matrix, as indicated by Figure 1A they are not major contributors to biofilm elasticity.

**Figure 2:**
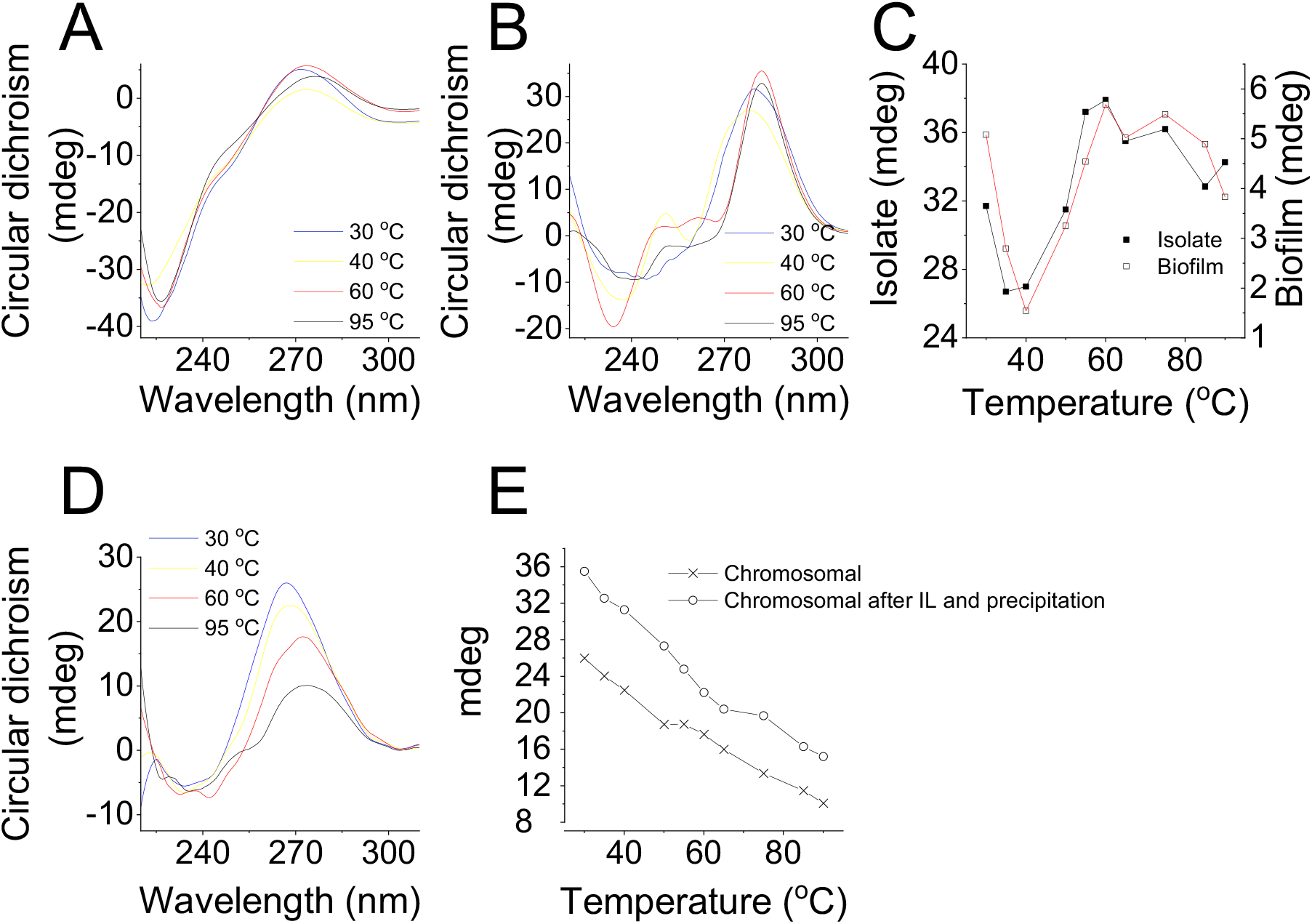
Representative circular dichroism (CD) spectra of (A) *Pseudomonas aeruginosa* biofilm and (B) eDNA gel isolate at 30°C (blue), 40°C (yellow), 60°C (red) and 95°C (black) dialysed against double distilled water. (C) Amplitude of dominant NA peak, CD_max_ (260-285 nm), from CD spectra of *P. aeruginosa* biofilm (seen in (A)) and its extracted extracellular nucleic acid gel (i.e. isolate) (seen in (B)) from T = 30°C to T = 95°C. (D) Representative CD spectra of *Pseudomonas aeruginosa* biofilm chromosomal DNA in double distilled water at 30°C (blue), 40°C (yellow), 60°C (red) and 95°C (black). (E) Amplitude of dominant NA peak, CD_max_ (260-285 nm), from CD spectra of *P. aeruginosa* biofilm chromosomal DNA (seen in (D)) from T = 30°C to T = 95°C with and without solubilization in 1-ethyl-3-methylimidazolium and fractional precipitation.

The CD spectrum of the isolated eDNA gel (Figure 2B) is dominated by the same NA peak that appears in the biofilm CD spectrum. CD melting experiments were then performed on both samples. The resulting melting curves (Figure 2C) show that their NA peak absorbance values follow the same trend across the temperature profiles with a maximum at their phase transition, or melting temperatures (i.e. T_m_) of (i.e. ≈ 57°C), indicating that melting occurs due to an eDNA gel-sol transition and that heating disrupts the higher order structure of NA. In contrast, the NA peak in chromosomal DNA extracted from *P. aeruginosa* planktonic cells decayed steadily with temperature (Figure 2D and 2E). This was also observed for chromosomal DNA processed through ionic liquid (i.e. EMIM-Ac dissolution and fractional precipitation).

Thus, *P. aeruginosa* eDNA has a different higher order structure to its chromosomal DNA. The elastic behavior observed throughout purification and faithfully preserved in the isolated eDNA, is not a consequence of the processing method. The extracted eDNA gel isolate is therefore an accurate proxy for understanding the intermolecular interactions that stabilize the extracellular matrices of *P. aeruginosa* biofilms. Moreover, the CD spectrum of the eDNA gel (Figure 2B) indicates multiple nucleic acid conformations. In addition to the peak at 272 nm, there is a trough that shifts from 245 to 230 nm with increasing temperature, another maximum at 215 nm that remains constant with temperature, and other sub-maxima at 260 and 253 nm that change with temperature. The absence of a trough at 200-215 nm precludes the possibility of NA in A- or Z-conformations (31). The dominant maximum and minimum could indicate either B-DNA or G-quadruplex conformations (33), although the peak at 215 nm is not a feature of B-DNA CD spectra and is slightly higher than the characteristic G-quadruplex low wavelength peak of 210 nm. Nonetheless, the appearance of several peaks in the NA region of 250-285 nm indicates that, while the dominant NA conformation is unclear, several conformations likely contribute to phase separation of the NA.

### Non-canonical base pairs or tetrads support extracellular network

Sequence analysis of the extracted material indicated that the gene coverage was even for both chromosomal and eDNA with the exception of bacteriophage Pf4 genes (Supplementary Figure 5). However, the Pf4 knockout mutant of *P. aeruginosa* also displayed an elastic response when dissolved in EMIM-Ac (Supplementary Figure 6, Supplementary Tables 1 and 2), indicating that Pf4 DNA is unlikely to be responsible for the phase-separating behavior of *P. aeruginosa* biofilms. The biophysical properties of eDNA cannot be explained by uneven gene coverage, including the expression of Pf4.

To describe the nature of nucleic acid base-pair interactions contributing to the elastic behavior of the isolated eDNA we performed Magic-angle spinning (MAS) solid-state NMR (SSNMR). This technique is ideal for intractable systems such as biofilms to eliminate solubility and extraction biases (34), and to analyze inter-molecular H-bond interactions to describe, for instance, inter-nucleotide base pairing in RNA (35). MAS SSNMR averages anisotropic interactions to provide high-resolution spectral characterization of insoluble and large biomolecular systems. By analyzing dipolar interactions, through-space heteronuclear correlations (e.g. N···H) can be detected.

To elucidate the mechanism of DNA gelation, we generated, by SSNMR, a 2D, through-space, ^15^N-^1^H HETCOR spectrum of ^15^N labeled eDNA gel isolated from *P. aeruginosa* biofilm matrix. The absence of signature protein and polysaccharide peaks in the ^15^N-^1^H HETCOR spectrum confirmed the complete absence of proteins and polysaccharides in the gel isolate (Supplementary Figure 7), which supports our assertions that neither contribute to eDNA gelation, despite the fact that both have been reported previously to co-localize with eDNA (17,21). The HETCOR spectrum across the imino proton region showed four signal clusters at *δH* 10-14 ppm, and *δN* 140-160 ppm (Figure 3), which arose from direct N-H couplings of T/U and G nucleobase imino groups. Two of these clusters (*δH* 12-14 ppm) resulted from imino protons hydrogen-bonded to a nucleobase nitrogen (i.e. N-H···N), and the other two (*δH* 10 - 12 ppm) from imino protons hydrogen-bonded to a nucleobase carbonyl oxygen (i.e. N-H···O) (36). Due to the longer cross polarization times we were able to observe a strong, long-range and indirect (i.e. intermolecular) correlation at *δN* 196 ppm arising from G-C Watson-Crick base pairing, i.e. C(N3)-G(H1). There was also a weak and indirect correlation at *δN* 220 ppm resulting from A-T/U Watson-Crick base pairing, i.e. A(N1) to U/T(H1).

**Figure 3:**
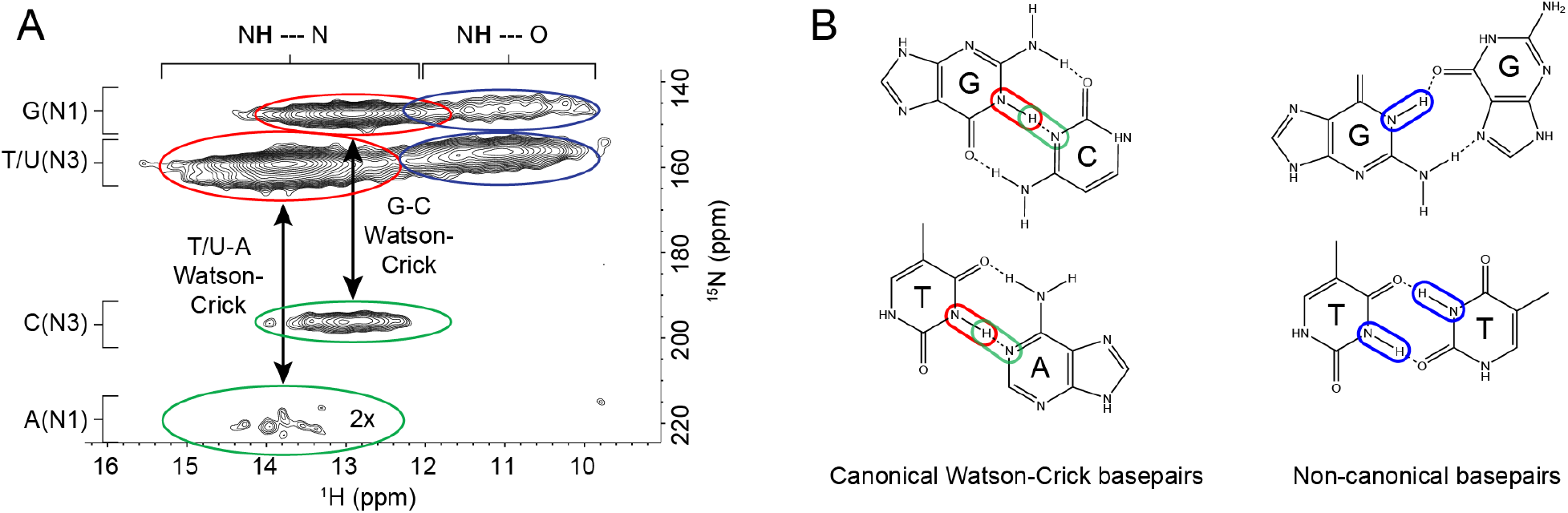
(A) Representative solid-state 2D ^1^H-^15^N through-space heteronuclear correlation (HETCOR) spectrum of extracellular nucleic acid (NA) gel isolate dialysed against double distilled water (2 mg), T = 25°C. Red ovals show the cross-peaks from direct N-H couplings of G (N1-H1) and T/U (N3-H3) when the protons were hydrogen bonded to nucleobase nitrogen atoms (i.e. NH···N). Green ovals present the cross-peaks generated form indirect (via hydrogen bonds) N-H couplings between the G(H1) and C(N3) as well as between T/U(H3) and A(N1). The threshold for the A(N1)-T/U(H3) correlation was increased due to lower signal intensity for that particular coupling. Cross-peaks denoted by red and green ovals indicate Watson-Crick G-C and U/T-A base pairings respectively. Blue ovals show cross-peaks from direct N-H couplings of G (N1-H1) and T/U (N3-H3) when the protons were hydrogen bonded to carbonyl oxygen (i.e. NH···O). These cross-peaks reveal non-canonical base pairings between G, T or U and G, T, U or C as well as the possibility of G-tetrad formations. (B) Canonical and non-canonical base pairing schematics illustrate the possible H-bond interactions between imino protons of A and G that can be observed in the HETCOR spectrum. The correlations are color-coded to match the spectrum. Covalent and hydrogen bonds are indicated by solid and dashed lines respectively.

The observation of NH to O interactions suggests the occurrence of non-canonical base pairs, triads or tetrads involving G and T/U. The absence of long-range correlations between the clusters at *δH* 10 - 12 ppm is also consistent with non-Watson-Crick pairings for the G and T/U nucleobases, where Hoogsteen H-bonded G-bases can assemble planarly into tetrads and self-stack to form structurally stable G-quadruplexes (37). *δH* assignments of imino protons H-bonded to nucleobase carbonyl O and N respectively are validated by 1-D ^1^H NMR spectra of well-characterized G-quadruplex and canonical B-DNA structures (Supplementary Figure 8A and 8B), and of a DNA quadruplex-duplex structural hybrid (Supplementary Figure 8C).

To determine whether non-canonical base pairs contribute to eDNA elasticity, a ^15^N-^1^H HETCOR spectrum was recorded for ^15^N labeled-eDNA gel after heating it to above the gel-sol transition temperature. In contrast to the sample without heating (Figure 4A), no non-canonical base-pair ^15^N-^1^H interactions were detected in the sample heated to 65°C (Figure 4B). The Watson-Crick base-pair interactions, on the other hand, persisted with and without heating. Similarly, the chromosomal DNA of *P. aeruginosa*, which does not exhibit the elastic or gel-like behavior characteristic of the eDNA (Figure 2D and 2E), displayed no NH to O interactions, which is consistent with the absence of non-canonical base-pair interactions (Figure 4C). Only canonical base-pair interactions were observed in the ^15^N-^1^H HETCOR solid state spectrum of P. aeruginosa PAO1 chromosomal DNA before (green) and after (gold) receiving the same treatment as eDNA, demonstrating that non-canonical base pair interactions are specific for the eDNA of *P. aeruginosa* and not an artifact of extraction and processing. Hence, the non-canonical base pairs, triads or tetrads are characteristic of eDNA in a gel or networked state and may contribute to stabilizing the higher order structure of eDNA networks.

**Figure 4:**
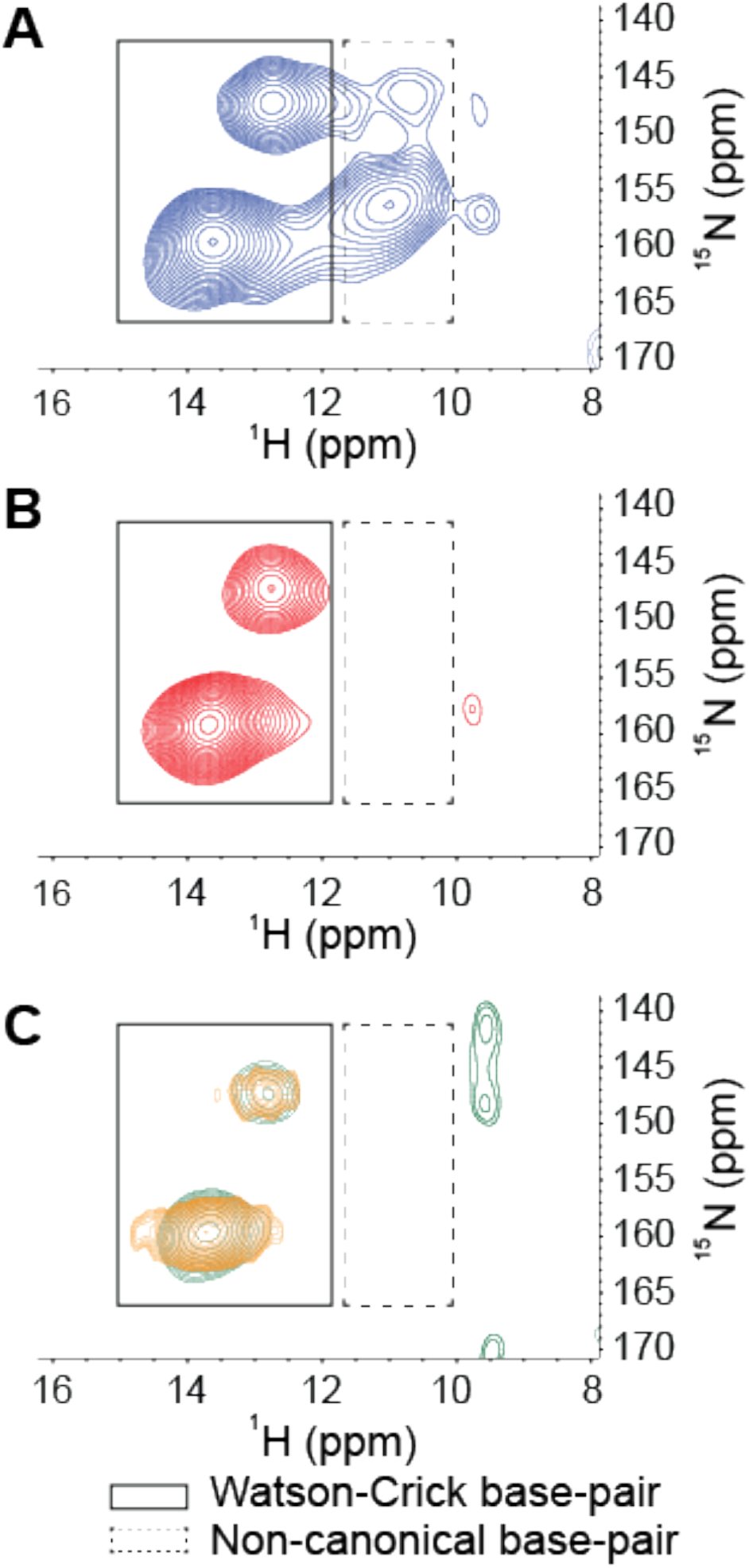
Representative solid-state 2D ^1^H-^15^N through-space heteronuclear correlation (HETCOR) spectrum of ^15^N-labelled extracellular nucleic acid (NA) gel isolate in double distilled water (2 mg), including direct N-H coupling region only, at T = 25°C with no pre-heating (A) and with pre-heating to 65°C (B), and ^15^N-labelled chromosomal DNA extracted from *Pseudomonas aeruginosa* biofilms in double distilled water before (green) and after (gold) receiving the same treatment as the extracellular nucleic acids (C), indicating the loss of non-canonical base pairings, triads or tetrads in the gel isolate upon heating and their absence in chromosomal DNA.

### Purine-rich RNA is also a component of eDNA gel

Total correlation spectroscopy (TOCSY) and ^1^H-^13^C heteronuclear single quantum coherence (HSQC) are nuclear magnetic resonance (NMR) techniques that can identify proton NMR correlations within individual ribose sugars and their proton-carbon single bond correlations respectively. Raising the pH can solubilize *P. aeruginosa* biofilms (38) and here we show that after alkalinization, the NA peaks dominate the solution ^1^H-^13^C HSQC spectra for the isolated material which provides further evidence that proteins or polysaccharides were absent (Supplementary Figure 9A) (39). However, the ^1^H-^13^C HSQC-TOCSY spectrum of the isolated gel when dissolved by alkalinization showed two clusters of sugar proton peaks (C1’-H1’) with correlations to neighboring carbons (C2’-H1’) (Figure 5A). This demonstrates that there are two types of nucleic acid present and contributing to the composition of the isolated gel.

**Figure 5:**
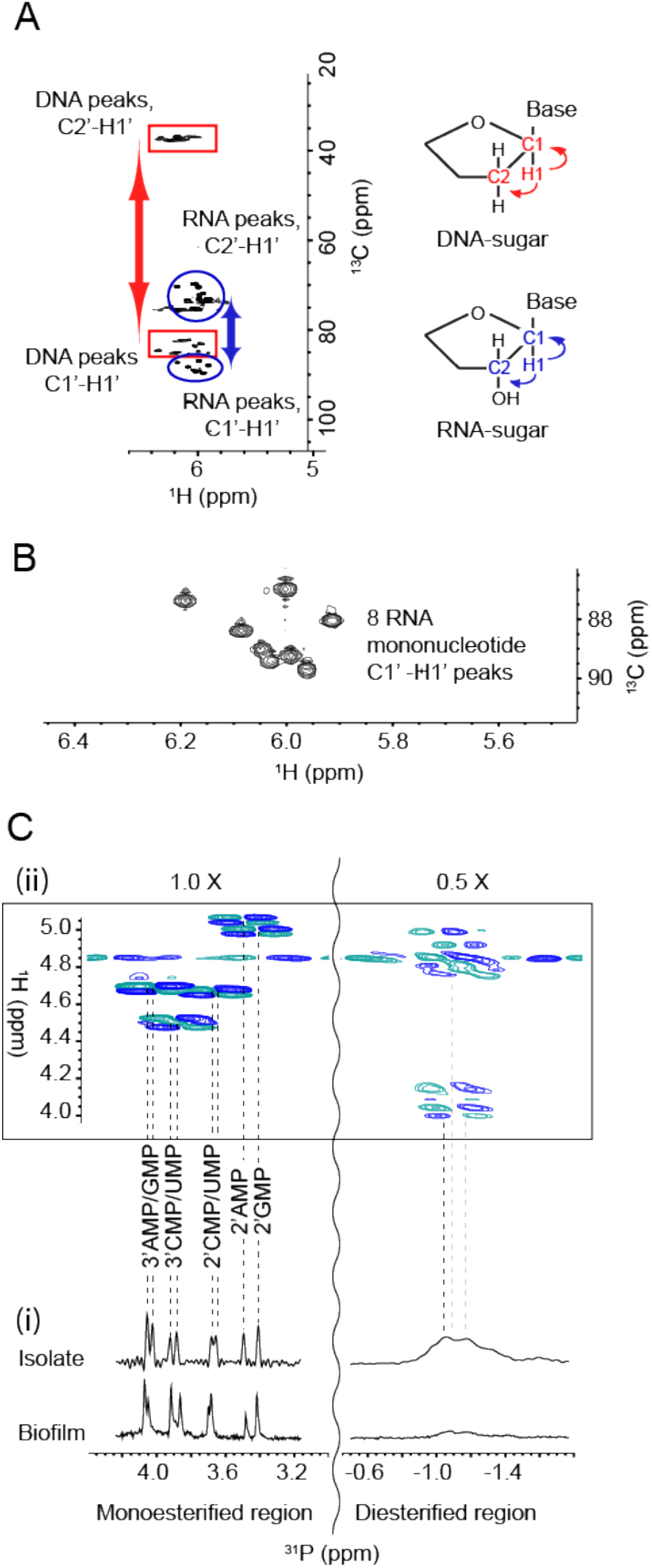
(A) Representative Nuclear magnetic resonance (NMR) ^1^H-^13^C heteronuclear single quantum coherence (HSQC)-total correlation spectroscopy (TOCSY) spectrum at 25°C of extracellular nucleic acids (NA) gel isolate after alkalinization showing the C1’-H1’ cross peaks of RNA (blue ovals) and DNA (red rectangles) and their correlations to the neighboring carbon C2’-H1’. Schematics (right) illustrate these correlations. (B) ^1^H-^13^C HSQC spectrum of extracellular eDNA gel isolate at 25°C after alkalinization identifying eight monoribonucleic acid ribose spin systems. (C) 1-D ^31^P NMR of NA isolate gel and *Pseudomomonas aeruginosa* biofilm with proton decoupling showing the presence of monoesterified (i.e. monoribonucleotides) and diesterified (i.e. DNA) phosphate peaks (i), and 2-D ^1^H-^31^P heteronuclear correlation (HETCOR) spectrum of extracellular NA showing the ^31^P-^1^H cross-peaks of monoribonucleotides and DNA (ii). Couplings of monoesterified phosphates to H2’ or H3’ of eight monoesterified monoribonucleotides (from left to right: 3’ AMP/3’ GMP, 3’ CMP/3’ UMP, 2’ CMP/2’ UMP, 2’ AMP, 2’ GMP); and diesterified phosphate to DNA H3’ and H5’/H5’’ protons, are denoted by the dashed lines. There is a discontinuity (wavy line) in the ^31^P axis due to the different thresholds required to illustrate the ^31^P-^1^H correlations in the mono-esterified and di-esterified regions. All samples were prepared in 0.1 M NaOD (10 mg/mL) and preheated to 55°C for 2 h.

The first cluster (rectangles) has C2’ chemical shift values of ~40 ppm and the second cluster (ovals) has C2’ values of ~70 ppm. The ^1^H-^13^C correlations denoted by the rectangles therefore arise from deoxyribose and those denoted by the ovals from ribose sugar conformations. The broadened form of the deoxyribonucleotide peaks is consistent with high molecular weight (MW) molecules, while the sharpness of the ribonucleotide peaks suggests that they arise from small nucleotides. Hence, eDNA and small ribonucleotides are both likely to be present in the gel dissolved by alkalinization.

Spin-systems (i.e. groupings of nuclei that do not couple with nuclei beyond their respective systems) for the eight sharp ribonucleotide peaks (Figure 5B) were assigned by HSQC-TOCSY and correlation spectroscopy (COSY) (Supplementary Figure 9B) with absolute identification achieved by heteronuclear multiple bond correlation spectroscopy (Supplementary Figure 10). The ^31^P NMR spectrum of the eDNA gel isolate at elevated pH revealed the presence of a mixture of monoesterified (3.4 to 4.1 ppm) and diesterified phosphates (−0.8 to −1.2 ppm), indicating the coexistence of monoribonucleotides and DNA respectively (Figure 5C (i)). Long-range ^31^P-^1^H correlations (i.e. from NA phosphorous to adjoined ribose proton) were observed in the ^31^P-^1^H heteronuclear correlation (HETCOR) spectrum (Figure 5C (ii)) for both monoribonucleotides and DNA. The broad ^31^P-^1^H cross peaks at ~-1.0 ppm (Figure 5C (ii)) correspond to long-range correlations between DNA phosphorous to H3’ and H5’/H5’’ protons. The eight monoribonucleotides were assigned to 2’- and 3’-(A, U, G, and C)-monophosphates. We therefore have both 2’ and 3’ monoribonucleotides present with the eDNA in the gel solubilized under alkaline conditions.

Importantly, the monoribonucleotide peaks only became resolved upon alkaline dissolution of the biofilm, in contrast to the broad peaks evident at pH 7 (Supplementary Figure 11A). This indicates that the monoribonucleotides are derived from chain structures that exist at biological pH, which is consistent with the observation in Figure 1A that RNA is a major contributor to *P. aeruginosa* biofilm elasticity.

The molar ratio of the individual ribonucleotides could be determined from the ^31^P spectrum of the gel dissolved at high pH (Table 1). While several peaks could not be separated, it was possible to deduce that the RNA is purine rich (i.e. 57 mol% A+G) and that the G+C mol% content of 46-50% differs from that of the *P. aeruginosa* genome (i.e. 67 mol%) (40). Quantification was undertaken on the basis of relative ^31^P NMR peak areas for the eight monoribonucleotides and is therefore unaffected by the presence of any free nucleic acid bases. The same peaks were also observed in the biofilm ^31^P spectrum after alkalinization (Figure 5C(i)). Sequence analysis of the eDNA (Supplementary Figure 5) indicated that, on the other hand, eDNA nucleobase ratios were consistent with the genome and the presence of canonical nucleotides.

In contrast to the liquid state ^31^P NMR spectrum of the alkali-dissolved isolate (Figure 5C (i)), the ^31^P SSNMR spectrum of the gel isolate (i.e. no alkali treatment; Figure 6A (i)) showed a single peak in the diesterified phosphate region, consistent with the presence of both DNA and RNA chains. There are no sharp peaks present in the monoesterified phosphate region, further demonstrating that the monoribonucleotides are a consequence of alkali transesterification (41).

**Figure 6:**
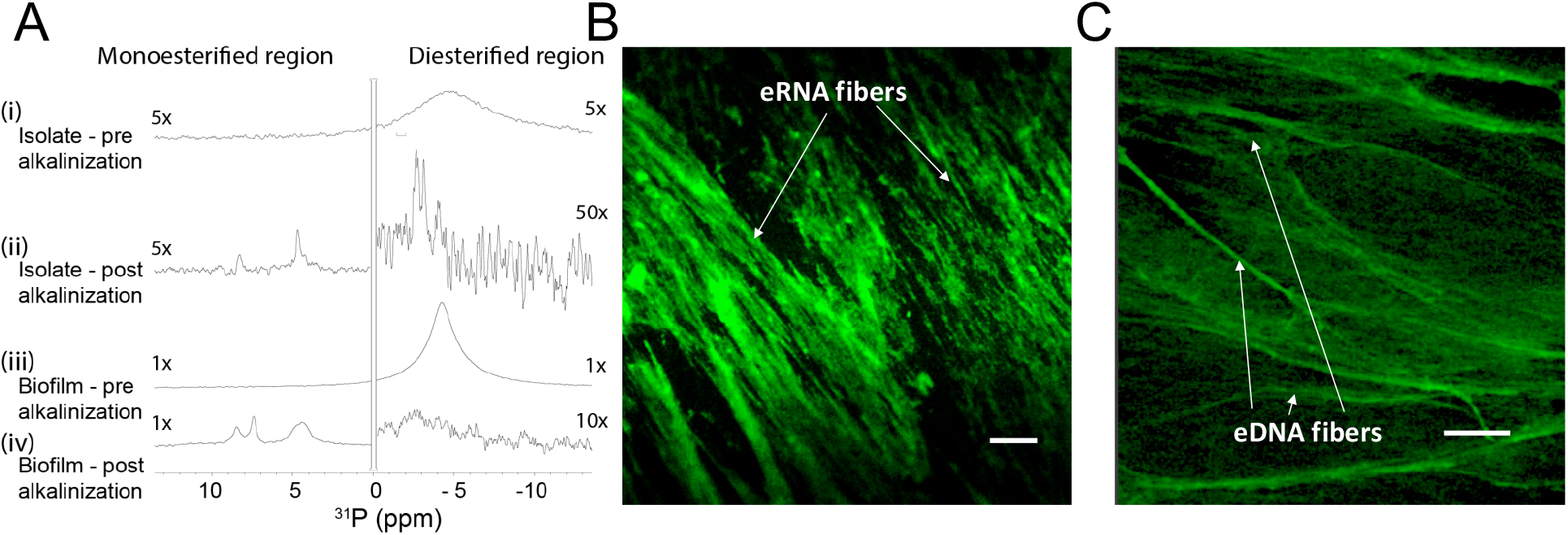
(A) Solid-state ^31^P NMR spectra at T = 25°C of extracellular eDNA gel isolate (i), alkalinized and lyophilized eDNA gel isolate (ii), *Pseudomonas aeruginosa* biofilm (iii) and alkalinized and lyophilized *P. aeruginosa* biofilm (iv) showing the presence of diesterified phosphate peaks and the absence of monoesterified phosphate peaks for both eDNA gel isolate and biofilm in double distilled water, and the coexistence of diesterified and monoesterified phosphate peaks for both samples after alkalinization. This indicates that alkalinization of the matrix results in RNA transesterification. Micrographs of *P. aeruginosa* eDNA gel isolate stained green with (B) SYTO RNASelect showing RNA fibers and (C) TOTO-1 showing DNA fibers (scale bar 10 μm).

Only the diesterified phosphate peak was observed in the ^31^P SSNMR spectrum of the biofilm (Figure 6A (iii)), while both RNA-derived monoesterified and DNA-derived diesterified phosphate peaks were present in the ^31^P SSNMR spectrum of alkali digested eDNA gel-isolate and biofilm (lyophilized) (Figure 6A (ii) and Figure 6A (iv), respectively). Complete alkali RNA transesterification was confirmed by the full conversion of diesterified to monoesterified phosphates (from liquid state ^31^P NMR spectra) for the RNA standard, when dissolved in 0.1 M NaOH (Supplementary Figure 11B). Conversely, RNA diesterified phosphate peaks were preserved in the RNA standard spectrum following dissolution in EMIM-Ac and recovery by perchloric acid (Supplementary Figure 11C). Hence, the alkaline conditions break down the RNA chains into individual monoribonucleotides, while the ionic liquid-based extraction does not. This suggests that the conventional method for *P. aeruginosa* biofilm dissolution in alkali (38) may in fact work by transesterifying RNA as a primary structural component and illustrates the importance of the ionic liquid-based extraction protocol described here as a non-destructive method for interrogating biofilm structural polymers.

To visualize the DNA-RNA network organization, we used nucleic acid-specific stains, i.e. RNA specific SYTO RNASelect dye (Figure 6B) and the eDNA-specific TOTO-1 dye (Figure 6C) (42). Fibrous structures, which are typical of networked polymer gels (43), were observed in the eDNA gel isolate stained with both nucleic acid dyes as well as in the biofilm stained with the eDNA specific dye (Figure 1D). This is consistent with our observation that the biofilm is readily degraded by DNase I (Figure 1A) (44). Furthermore, the eDNA gel isolate was stained positively with the RNA-specific dye even after DNase I digestion, providing further evidence that RNA contributes to networking (Supplementary Figure 12A). However, digestion with RNases III, A and H, did not degrade the biofilm (Figure 1A; Supplementary Figure 12B). This may suggest that the RNase binding was shielded by hairpins in the network, or the presence of non-canonical DNA-RNA interactions (45,46). Additionally, while eDNA gel isolate treated directly with RNAse A was not reduced in size (i.e. 1000–4000 nucleotides; Supplementary Figure 12C, Lane D1), gel isolate pre-digested with DNase I (Lane E1) and subsequently digested by RNase A (F1), was degraded from 350-1000 nucleotides into a single band of approximately 150 nucleotides. This provides further evidence that extracellular RNA, of MW 350 to 1000 nucleotides, are protected from RNase A digestion because they are likely complexed to eDNA. The ScreenTape analysis also shows the absence of clear 23S and 16S RNA bands, unlike the control (Lane G1), indicating that extracellular RNA are either partially degraded or not ribosomal RNA, and highlighting the challenges in detecting and analyzing RNA in the extracellular matrix.

The presence of purine-rich RNA was confirmed by LCMS analysis, where the eDNA gel was found to consist of about 4 % w/w RNA although the actual amount is likely to be higher due to incomplete transesterification. Nonetheless, the structural importance of RNA is illustrated by the irreversible pH dependent gel-sol transition displayed by the biofilm (Figure 1C), which occurs coincident with transesterification of RNA and not DNA (Figure 5). If the role of RNA is to facilitate cross-linking of DNA, relatively low amounts of RNA would be required to facilitate its cross-linking.

## Discussion

We extracted a gel-forming complex composed of DNA and RNA from the extracellular matrix of *P. aeruginosa* biofilms without disrupting its fundamental structural or chemical organization. Long-term stability has been reported for various biomolecules in ionic liquids, including DNA (47), and its use here for assessing higher-order eDNA structure is informative. We report the novel observation that extracellular RNA contributes structurally to biofilms and that it contributes along with eDNA to the fundamental viscoelasticity of *Pseudomonas* biofilms. A viscoelastic matrix differentiates planktonic cells from biofilms and the viscoelastic property of the matrix itself is considered a virulence factor in chronic biofilm infections (8). Our data suggest that in forming biofilms *Pseudomonas* assemble their RNA and DNA into an extracellular framework that supports viscoelasticity. Understanding how the DNA and RNA is organized extracellularly during different development stages and in response to environmental conditions is thus central to understanding *Pseudomonas* biofilms.

While gel-forming exopolymers have been detected in other biofilms (48), no functional DNA gel has been described for any biological system, much less one also including extracellular RNA. DNA hydrogels are attractive biomaterials due to their biocompatibility, however DNA gelation typically requires either ligase-mediated reactions to induce branched DNA molecule formation (i.e. Y-scaffold), chemical crosslinking agents (e.g. epoxides), cationic poly(electrolytes) (e.g. spermidine) to promote complexes by electrostatic interactions, or physical entanglements (49). The presence of RNA with eDNA therefore addresses the question of how DNA, otherwise favoring linear or circular structures, can form networked structures in biofilms in the absence of, for example, cationic polyelectrolytes. RNA is more structurally versatile than DNA (e.g. hairpins), has a greater tendency to form non-canonical base pairs, and has been shown to undergo a sol-gel transition due to multivalent interactions between strands (50). It informs on why *Pseudomonas* biofilms are readily dissolved by alkalinization, resulting from RNA transesterification, providing further evidence of the structural role of RNA. It also suggests RNA as potential targets for biofilm control, and there are several possible explanations for why RNA nucleobase ratios might deviate from genomic abundances (i.e. 67% G-C for *P. aeruginosa*) (51), including small-RNA, microRNA, non-coding RNA, virus RNA and alternative polyadenylation (52).

While it is not possible to identify the precise nature of the DNA-RNA interaction based on the information presented in this study, purine-rich RNA as described here for the extracellular eDNA gel isolate (57 mol% A-G) is a characteristic of DNA-RNA duplexes because RNA purines bind more strongly to their DNA pyrimidine complement than vice versa (53). Both Watson-Crick and non-canonical interactions were present in the eDNA gel isolate, which is also consistent with the discrepancy observed between RNA and DNA G+C mol% contents (Table 1). It is possible, therefore, that interactions between DNA and RNA, as described in this study, account for the formation of a highly stable nucleic acid gel.

The results presented provide unprecedented resolution of the biofilm exopolymeric matrix and its key foundation structural components. We have developed a methodology that preserved the molecular organization of the foundational polymer in its native state upon extraction and isolation. This enabled us to use SSNMR to describe intermolecular associations at the atomic level.

This study demonstrates that an eDNA-RNA network, responsible for the elastic gel-like properties of the matrix, contributes greatly to the formation of a physically distinct habitat in the matrix of *P. aeruginosa* biofilms, which in turn is responsible for emergent properties such as nutrient accumulation, stress and antibiotic resistance, and shelter (54). This finding is a departure from the prevailing paradigm that biofilm gelation is only due to polysaccharides. The elastic and biofilm-forming properties were also observed for *Pseudomonas putida* and *Pseudomonas protegens* (Supplementary Figures 6 and 13; Supplementary Tables 1 and 2) suggesting that eDNA is broadly important for biofilm formation in member species of this genus. In addition to *Pseudomonas* spp., other organisms are known to have eDNA, including the clinically relevant organisms *Staphylococcus aureus* (55)*, Staphylococcus epidermidis* (56) and *Mycobacterium abscessus* (57) as well as environmental isolates such as strain F8 from the South Saskatchewan River (16).

While eDNA is commonly thought to be the product of cell lysis, and has been shown to be released by a sub-population of lytic *P. aeruginosa* cells (19), there is an increasing awareness that it serves an important structural function, such as in activated sludge granules (58). Additionally, Bockelmann et al.(16) observed that stable filamentous networks produced by the aquatic strain F8 were comprised of DNA. The present study provides the first account of a structural signature for eDNA, absent in the chromosomal DNA. However, the environmental factors contributing to eDNA release and how this is regulated (i.e. whether it is an active or passive process) are still unclear. It is possible that DNA-RNA foundational gel networks, as described here for the Pseudomonads, are broad-scoping phenomena in biofilms, and that interactions with RNA enable an extracellular structural function for DNA more broadly. Thus, elucidating how nucleic acids, including RNA, are integral to the biophysical and other emergent properties imparted on the biofilm via the extracellular matrix, will inform the regulation and control of extracellular nucleic acid release across environmental and clinical biofilms.

## Materials and Methods

### Bacterial strains

Unless otherwise stated, all experiments were undertaken directly on *P. aeruginosa* PAO1 biofilms grown in lysogeny broth (LB) at 37°C. The *P. aeruginosa* PAO1 Δ*pf4* knockout mutant is a defined Pf4 chromosomal deletion mutant of the entire Pf4 prophage genome (59). *P. aeruginosa* PDO300, PDO300 (*Δpel*) and PDO300(*Δpsl*) mutant strains were gratefully received from Professor Bernd H. A. Rehm, Institute of Molecular Biosciences, Massey University, New Zealand (28), where PDO300 is an isogenic *mucA* deletion mutant of PAO1 that overproduces alginate. PDO300, PDO300 (*Δpel*), PDO300(*Δpsl*) and PAO1 (*Δpf4),* were also grown in LB at 37°C. *P. protogens Pf-5* and *P. putida* ATCC BAA-477 and S12 strain were grown in LB at 30°C.

### Biofilm growth assay

Ten milliliter aliquots of *P. aeruginosa* planktonic pre-cultures (LB, 200 rpm, 37°C, OD_600_ 2.40, 16 h) were diluted 50 times with LB in 2 L Erlenmeyer flasks and incubated for 5 d under static conditions. Supplementary Figure 14A displays an image of 5 d old biofilm in LB. The cultures were centrifuged at 10,000 × *g* for 15 min, the supernatant removed by decanting, and the biofilm then collected and lyophilized (LabConco). Supplementary Figure 14B displays an image of the culture after centrifugation showing clear separation of biofilm and supernatant.

### Enzymatic digestions

Twenty milligrams of lyophilized biofilm was resuspended in 1 mL of either i) RNase buffer (50 mM Tris HCl, 10 mM EDTA, pH8) with 0.2 mg RNaseA from bovine pancreas (Sigma Aldrich), ii) storage buffer (10 mM NaCl, 10 mM Tris-HCl) with 0.1 mg Pronase E from *Streptomyces grisens* (Sigma Aldrich) with 0.5% (v/v), iii) 1X RNase H reaction buffer (20 mM Tris-HCl pH 7.8, 40 mM KCl, 8 mM MgCl_2_, 1 mM DTT) with 0.4 mg RNase H (Thermo Fisher Scientific), iv) 1X RNase III reaction buffer (500 mM NaCl, 100 mM Tris pH 7.9, 100 mM MgCl_2_, 10 mM DTT) with 0.6 mg RNase II (Thermo Fisher Scientific) or v) DNase I buffer (100 mM Tris (pH 7.5), 25 mM MgCl_2_ and CaCl_2_) with 0.2 mg DNAse I (active and inactivated) from bovine pancreas (Sigma Aldrich). DNase I was inactivated with the addition of 1 mM EDTA and following heating to 85°C in DNase I buffer for 10 min. All digestions were performed with shaking at 200 rpm at 37°C for 16 h. The suspensions were then centrifuged (10,000 × *g*, 15 min), the supernatant was discarded and the pellets of the biofilm materials were lyophilized.

### Normal force measurement

Forty milligrams per milliliter solutions of lyophilized biofilms were added to 1 mL 1-ethyl-3-methylimidazolium acetate and incubated at 55°C for 2 h. A Haake Mars 3 (Thermo Fisher Scientific) stress-controlled rotational rheometer with Peltier controlled element at 25°C, was used for rheological measurements. Thirty five-millimeter diameter parallel plate geometry was used with smooth titanium plates to measure viscosity and normal stress difference (*N*_1_ – *N*_2_) where *N*_1_ is the difference between normal stresses in the direction of shearing and those oriented perpendicular to the shear plane, and *N_2_* the difference between normal stresses perpendicular to the shear plane and those in the neutral, traverse, direction (26). Prior to measurement, the gap error was zeroed at 4 N and gap error calculated as previously described (60–62). The parallel plate rheometer was used because it allows rheological characterization of ultra-low sample volumes and permits access to high shear rates for characterization of non-linear elasticity without measurement artifacts such as inertia (i.e. Reynolds number is low). One hundred microliters of sample were deposited on the plates. The plates were closed to 100 μm, the sample trimmed and the sample allowed to sit for 5 min prior to measurement. All measurements with Normal force (*F_N_*) less than the resolution of the rheometer (i.e. < 0.1 N) were set to 0 before calculation of *N_1_*-*N_2_* using equations 9-11, and viscosity from equation 3-5 in Davies and Stokes (60) for the parallel plate geometry.

Only the linearly increasing portion of the normal stress difference curves are presented. Above this range normal stress difference begins to decrease again which may be due to elastic instabilities or associating polymers (63). Corrections were made to *N*_1_ – *N*_2_ to account for inertia using equation 17 in Davies and Stokes (60) and to correct for the baseline residual force in the samples. Except for the DNase I-treated biofilm, the shear rheology for all treatments could be modelled using the finitely extensible non-linear elastic with Gaussian closure proposed by Peterlin (FENE-P) constitutive model by varying four parameters to fit shear viscosity and normal stress difference as a function of shear rate (Supplementary Table 2). To use this model, we make the assumption that *N*_2_ << *N*_1_, and thus ΔN ~ N_1_, as is common for polymer solutions (Davies and Stokes (60)). Fitting parameters for the FENE-P model include *λ*_1_ = relaxation time, b = a measure of the relative extensibility of the model spring, *η*_s_ = solvent viscosity, *η*_p_ = polymer contribution to the viscosity. The FENE-P equations can be written in the following format, as shown by Bird et al.(61):

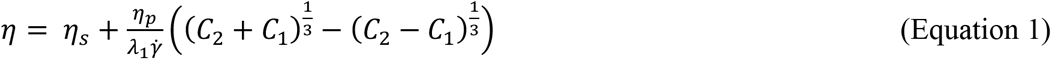

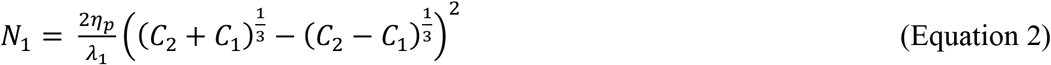

Where:

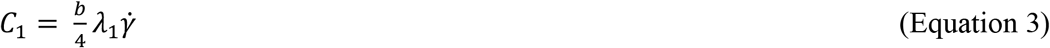

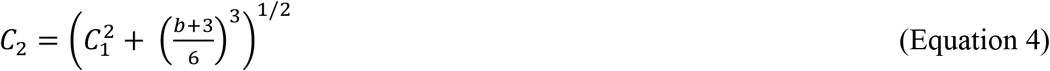

All measurements were performed in triplicate. For clarity, one representative data set is presented in Figure 1A-B and Supplementary Figure 1A-B, with the respective Power Law and FENE-P model fit to that data set. Averaged values for the FENE-P and Power law fits, with the standard deviation across three biological replicates are shown in Supplementary Tables 1 and 2, where each biological replicate refers to a separate growth assay.

For oscillatory measurements a Haake Mars 60 (Thermo Scientific) stress-controlled rotational rheometer with Peltier controlled element at 25°C was used. 0.25 mL biofilm (native wild type, wild type following digestion with active and inactivated DNase I, RNase III and RNase H, PDO300(*Δpel*) and PDO300(*Δpsl*) and eDNA gel were collected from solution and deposited on thirty five-millimeter diameter parallel plate geometry with smooth titanium plates, which were closed to 250 μm, and the sample trimmed prior to measurement. A frequency range of 0.1 to 1 s^−1^ was used and at a strain in the linear viscoelastic region for each of the samples analyzed.

### Extracellular polymeric substances (EPS) extraction

Lyophilized biofilms were dissolved in ionic liquid mixture (40% (v/v) 1-ethyl-3-methylimidazolium acetate (EMIM Ac): 60% (v/v) N,N-dimethyl acetamide (DMAc)) at 55°C for 16 h. The solution was centrifuged (10,000 × *g*) to remove any undissolved material. Perchloric acid (70%) was added (0.05% v/v) to the viscosified centrate (on ice). After 15 min incubation, the solution was centrifuged at 10,000 × *g* at 4°C for 15 min and the pellet recovered. This was repeated on the centrate two to four times until the solution was not viscous. The precipitate was dialysed against double distilled water for 2 d at 4°C (SnakeSkin™ Dialysis Tubing, 3.5K MWCO, 22 mm) and the retentate lyophilized (FreeZone Plus 4.5 Liter Cascade Benchtop Freeze Dry System). The same procedure was performed in triplicate on calf thymus DNA, lipase, cytochrome C for the purposes of determining recovery yield of representative exoproteins, and on RNA from torula yeast for assessing by ^31^P NMR whether the extraction procedure contributed to RNA transesterification (all from Sigma Aldrich).

### Extracellular nucleic acid isolation

Twenty milligrams of lyophilized retentate (i.e. post perchloric acid precipitation) were dissolved in 1 mL of 40% (v/v) EMIM-Ac, 60% DMAc (v/v) (55°C, 16 h). Chromatographic separation was achieved in a Shimadzu system comprising DGU-20A 3r Prominence Degasser and LC-20AD Solvent Delivery Unit, fitted with two Agilent PLgel 10 μm column of 10^5^Å pore size for separation across the MW range 200 kDa to 2000 kDa. The eluent flow rate was 3.0 mL.min^−1^ and the injection volume 1 mL. The fractions with molecular weight range of 2000-800kDa and 800-200kDa were pooled, precipitated with 60% v/v ethanol, and the precipitate resuspended in doubled distilled water and dialyzed for 2 d at 4°C (SnakeSkin™ Dialysis Tubing, 3.5K MWCO, 22 mm) against double distilled water to induce gelation. The gel was then collected from the dialysis tubing.

### eDNA temperature-dependent phase transition

Five-day old biofilm and eDNA gel isolate were resuspended in double distilled water to achieve UV absorbance reading 1 and ddH_2_O served as a blank. The heat-treated samples were analyzed by JASCO-815 spectropolarimeter in a 1 cm path length quartz cuvette containing a solution volume 500 μL. Spectra (200-320 nm) were measured at 1°C increments from 30 - 95°C. For each measurement, an average of three scans was taken and the buffer spectra subtracted. The magnitude of the DNA peak presented is the rolling average across five temperatures. The differential scanning calorimetry thermogram was measured in a MicroSC multicell calorimeter (Setaram,), using Calisto for data collection and processing. 400 and 60 mg of 5 d biofilm and eDNA gel isolates respectively added to sealed-1 mL Hastelloy C cells for measurement. An empty HastelloyC cell was used as reference. The samples were heated from 20-100°C at 1 K.min^−1^, then cooled down to 20°C at 1 K.min^−1^. The experiment was run under nitrogen gas sweeping.

### Solid-state NMR

For solid-state NMR experiments performed on the eDNA gel isolate, ^15^N labeled NH_4_Cl-supplemented M9 minimal media was used for biofilm growth. M9 consisted of 9.552 g.L^−1^ Na_2_HPO_4_.2H_2_O, 4.41 g litre^−1^ KH_2_PO_4,_ 1.71 g.L^−1^ NaCl, 1 g.L^−1^ ^15^NH_4_Cl, 0.24 g.L^−1^ MgSO_4_, 0.011 g.L^−1^ CaCl_2_, 2 g.L^−1^ casamino acids, and 0.4 g.L^−1^ glucose. The eDNA gel was prepared from the M9 grown *P. aeruginosa* PAO1 biofilm as described above.

Solid-state NMR experiments were performed on 1) eDNA gel with and 2) without preheating to 65°C, and 3) chromosomal DNA after lyophilization, dissolution in (EMIM Ac): 60% (v/v) N,N-dimethyl acetamide (DMAc)) at 55°C for 16 h, precipitation with ethanol (60% v/v) and dialysis against water (3500 Da MWCO) as described above for eDNA. A 14.1 T Bruker Advance III instruments was used equipped with a 1.9 mm MAS probe operated in double mode. The typical ^1^H, ^15^N and ^31^P π/2 pulse lengths were 2.3, 3.7, and 4.5 μs, respectively. 2D dipolar-based ^15^N-^1^H heteronuclear-correlation (HETCOR) experiments were conducted on the ^15^N-labelled eDNA gel isolate at 37 kHz MAS spinning frequency. Variable temperature was regulated at −20°C and the sample temperature was 12°C (calibrated using ethylene glycol). In the ^15^N-^1^H HETCOR experiments, the initially excited ^1^H magnetization was transferred to ^15^N through a cross polarization step followed by t1 evolution. Then, the ^15^N magnetization was flipped to longitudinal axis and 400 ms proton saturation pulses were applied for water suppression. Subsequently, the ^15^N magnetization was flipped to the transverse plane and transferred to ^1^H via a second CP step for signal acquisition. Two ^15^N-^1^H HETCOR experiments were collected, one with 400 μs and the other with 2 ms contact times applied for both of the CP steps. Low power XiX ^1^H decoupling (~10 kHz) was employed during ^15^N evolution and WALTZ-16 decoupling (10 kHz) was implemented on ^15^N channel during ^1^H acquisition. 1-D ^31^P experiments were performed on ^15^N-labelled eDNA gel isolate and ^15^N-labelled *P. aeruginosa* PAO biofilm, both directly after dialysis against double distilled water at 4°C for 2 d (SnakeSkin™ Dialysis Tubing, 3.5K MWCO, 22 mm) and following alkalinization (0.1 M NaOD, 55°C, 15 min) and lyophilization (FreeZone Plus 4.5 Liter Cascade Benchtop Freeze Dry System). 15 kHz MAS spinning frequency and a sample temperature of 27°C. 75 kHz SPINAL64 ^1^H decoupling was applied during ^31^P acquisition time. All chemical shifts were indirectly referenced using adamantane as a secondary standard (downfield peak is at 40.48 ppm, DSS scale).

### Solution-state nuclear magnetic resonance (NMR)

Solution-state NMR experiments were performed on an 800 MHz Bruker Avance III spectrometer at 25°C. Sample concentration was 10 mg (dry weight).mL^−1^ unless otherwise specified. Spectra were recorded either under conditions of neutral pH in 100% D_2_O (Cambridge Isotope Laboratories), or following alkalinization (i.e. transesterification, 0.1 M NaOD, 55°C, 2 h). 1-D NMR experiments include ^1^H and ^31^P direct detection, while 2-D NMR analysis include ^13^C-HSQC, ^13^C-HSQC-TOCSY, ^1^H-^31^P HETCOR, HMBC, and COSY. All spectra were referenced internally against the solvent peak (90% H2O and 10% D2O, unless otherwise specified), and all spectral analyses were performed using Topspin 4.0 and NMRFAM-SPARKY (64) software.

Asolectin (Sigma Aldrich) standard (10 mg/mL) and lyophilized *P. aeruginosa* PAO1 biofilm (10 mg/mL) were dissolved in 40% (v/v) EMIM Ac: 60% (v/v) DMAc at (55°C, 2 h). *P. aeruginosa* PAO1 pre-culture cell lysate was prepared by lysing pre-culture cells with lysozyme in PBS. 10% (v/v) of D_2_O was added to all samples for locking purposes.

### Chromosomal DNA extraction

Chromosomal DNA was extracted from the biofilm using FastDNA SPIN Kit for soil (MP Biomedicals, USA) as per the standard protocol. Briefly, biofilm was resuspended in Sodium Phosphate Buffer in was lysed (Lysing Matrix), homogenized (FastPrep^®^, 40 seconds, speed setting 6.0), and the cell debris removed by centrifugation (14,000 × *g*, 5 min). Proteins were removed by precipitation (250 μl Protein Precipitation Solution), the supernatant mixed with DNA Binding Matrix, which was then homogenized and transferred to a SPIN™ Filter. Excess supernatant was removed by centrifugation (14,000 × *g*, 5 min). DNA was then eluted from air dried DNA Binding Matrix with DNase/ Pyrogen-Free Water.

### Nucleotide sequencing, size and compositional analysis

The eDNA gel isolate was resuspended in 500 uL of 1x Protease K solution (10× Protease K solution: 10 mM Tris HCl, 1% SDS and 10 mM EDTA, pH 8 buffer (10× protease K buffer containing 500 mM Tris-HCl, 10% SDS, 10 mM CaCl_2_) and 10 μL of Protease K (20 mg.mL^−1^, Thermo Fisher Scientific) was added and the mixture incubated at 60°C for 2-16 h, after which DNA was extracted as previously described in the phenol-chloroform method (65). Samples before sequencing were further purified to remove any remaining protein and RNA by RNase and Proteinase K treatment. The DNA was then isolated using phenol-chloroform precipitation as described above. The DNA precipitate was dissolved in TE buffer, the purity confirmed by 260/280 value in Nanodrop (acceptable range value: 1.8-2.0) and Qubit^®^ 2.0 fluorometer.

The molecular weight distributions of extracellular and genomic DNA were measured on a 1% agarose gel, which was prepared from Viviantis LE grade agarose using 1x TAE buffer (40 mM Tris, 20 mM Acetate and 1 mM EDTA, pH 8.6). Gels were run horizontally. After electrophoresis, the gel was stained for 0.5 h with ethidium bromide and visualized under UV(66).

Three biological replicates were used for each DNA sequence analysis. Library was produced using Illumina DNA sample preparation kit. The libraries were sequenced using Illumina MiSeq platform (Illumina, San Diego, Ca) with paired-end protocol to read lengths of 600 nt generating a total of 1,614,106 and 1,848,846 paired end reads. Raw reads were quality filtered (reads remaining after trimming: PPG1-1549104, PBLC1-1666280) and aligned to the *P. aeruginosa* PAO1 (AE004091) genome using CLC Genomics Workbench 9.0 (CLC bio, Cambridge, MA). Extracellular RNA length was determined by TapeStation, model 2200 (Agilent Technologies). Monoribonucleotide quantification was achieved on three biological replicates of the nucleic acid isolate gels, pre-alkalinized with 0.4 M NaOH in double distilled water (10 mg dry weight gel/mL) at 55 °C for 2 h. Extracellular monoribonucleotides resulting from transesterification were quantified using Waters ACQUITY UPLC™ (Waters Milford, MA) liquid chromatography system with XEVO TQ-S mass spectrometer (Waters Milford, MA) equipped with an electrospray ionization (ESI) source. 2 μl of diluted samples were injected for analysis. An AQUITY UPLC HSS T3 column (2.1 mm × 150 mm, 1.7 μm, Waters, USA) was used, maintained at 45°C. 0.1% v/v formic acid and acetonitrile were used to establish the required gradient. The elution gradient increased from 0-1% v/v acetonitrile in first 6 min and then to 100% at 7 min, which was maintained for another 1.5 min, reduced to 0% at 8.6 min and held until 12 min. Torula yeast RNA (0.5 mg/L) was used as RNA standard (Sigma Aldrich). The HPLC flow rate was maintained at 0.3 ml.min^−1^.

### Staining and microscopy

Microscopic imaging was conducted on a confocal microscope Zeiss LSM 780 with a 63× objective. Extracellular RNA in the gel isolate were stained using SYTO RNASelect (Thermo Fisher Scientific) green fluorescent cell stain (5 mM solution in DMSO). Five μM stain solution was prepared from 1 μL of stock in 1X PBS solution (137 mM NaCl,2.7 mM KCl,4.3 mM Na_2_HPO_4_,1.47 mM KH_2_PO_4_). The gel isolate was labelled with 5 μM stain solution, kept at 37°C for 20 min and then transferred to glass side for imaging. eDNA staining was achieved by depositing biofilm or eDNA gel isolate on a glass slide, air-drying overnight and incubating with 2 μM TOTO-1 iodide (1 mM solution in DMSO, Thermo Fisher Scientific) for 15 min.

## Acknowledgments

We acknowledge Prof Bernd Rehm for supplying polysaccharide deletion mutants of *P. aeruginosa*, Dan Roizman for providing *P. putida,* Long Yu for assistance with rheology, Dr Gleb Yakubov for coordinating sample preparation for rheological measurements, Ravi Jagadeeshan for discussion regarding DNA normal force analysis, and Florentin Constancias for analyzing the sequencing data. SCELSE is funded by Singapore’s Ministry of Education, National Research Foundation, Nanyang Technological University (NTU), and National University of Singapore (NUS) and hosted by NTU in partnership with NUS. J.R.S acknowledges the assistance of Australian Research Council Discovery Project DP180101919.

## Author Contributions

T.S., F.R.W., W.L.L., S.M., H.M.S., S.S.A. and X.S. performed experiments; T.S. and A.T.P. designed the experiments. T.S. (all), F.R.W. (nucleic acids), A.T.P. (nucleic acids), H.M.S. (rheology), G.S.K. (sequencing), S.S.A. (LCMS/MS) and J.R.S. (rheology) advised on measurements and analysed the data. T.S., F.R.W., W.L.L., H.M.S., S.A.R., A.T.P and S.K. wrote the manuscript. J.R.S edited the manuscript.

## Competing Interests

All authors have no competing interests.

## Notes

### Competing Interest Statement

The authors have declared no competing interest.

### Summary of Updates

Solid state NMR analysis of chromosomal DNA was undertaken with and without preheating to 60 degrees, and on the reconstituted extracellular DNA after preheating to 60 degrees, to demonstrate that the non-canonical base pair interactions exist only in the extracellular DNA (i.e. not chromosomal DNA) and only when it is in gel state (i.e. not after preheating to 60 degrees), further supporting our assertion that the non canonical base pair interactions contribute to eDNA gelation.

