## Supporting information for "The biofilm matrix scaffold of *Pseudomonas* species contains non-canonically base paired extracellular DNA and RNA"

^2^ School of Physical and Mathematical Sciences, Nanyang Technological University, 637371, Singapore.^3^ Materials and Chemical Characterisation Facility MC^2^, University of Bath, BA27AY, Bath, United Kingdom. ^4^ Singapore Phenome Centre, Nanyang Technological University, 636921, Singapore. ^5^ School of Chemical Engineering, The University of Queensland, 4072, Brisbane, Australia. ^6^ The iThree Institute, The University of Technology Sydney, Sydney, 2007, Australia. ^7^ School of Biological Sciences, Nanyang Technological University, 637551, Singapore. ^8^ Centre for Marine Science and Innovation, School of Biological, Earth and Environmental Sciences, University of New South Wales, Sydney, 2052, Australia.

**This PDF file includes:**

Supplementary Figs. 1 to 13

Supplementary Tables 1-2

| 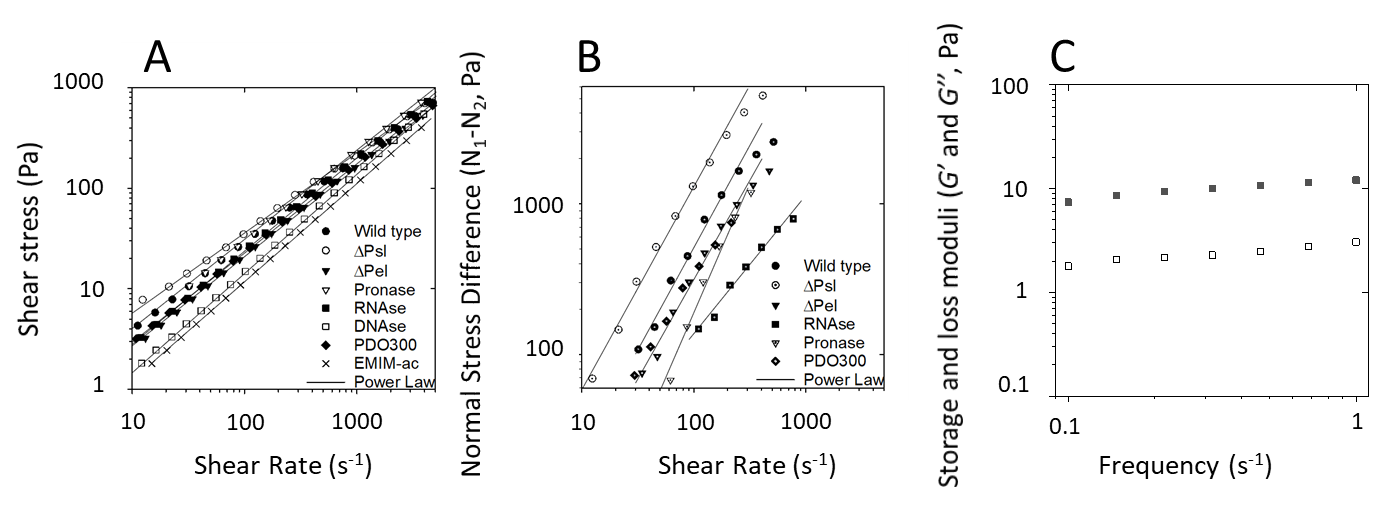 |
| --- |

**Supplementary Figure 1:** (A) Shear stress and (B) normal stress differences (*N_1_ –N_2_*) as a function of shear rate for *P. aeruginosa* biofilm: wild type; PDO300; ΔPsl, ΔPel; pronase digested; RNase A digested; and DNase I digested wild type biofilm, dissolved in 1-ethyl-3-methylimidazolium acetate (40 mg/mL) at 25 °C, 100 µm rheometer measurement gap, shear stress sweep from 10 to 1000 Pa. (*N_1_ –N_2_*) is not described for DNase I digested biofilm as its normal force (F_N_) is less than the resolution of the rheometer (i.e. 0.1 N) and is set to zero for calculating (*N_1_ –N_2_*). Lines indicate power-law fits to the data. The power law dependences of shear stress on shear rate (*m*) (see Table S1) indicate Newtonian-like rheological properties**.** (C) Frequency dependence of storage ($G^{'}$, closed shapes) and loss ($G^{''}$, open shapes) of *Pseudomonas aeruginosa* wild type biofilm digested by heat-inactivated DNase, at 25 °C (250 μm plate gap, 0.1% strain). Note, *G’* is not measurable in the DNase I treated biofilm.

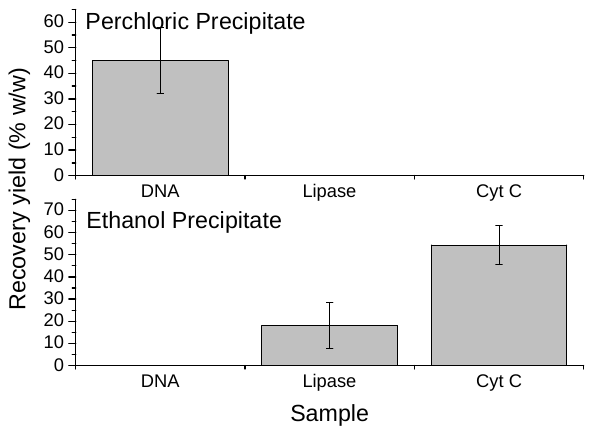

B

A

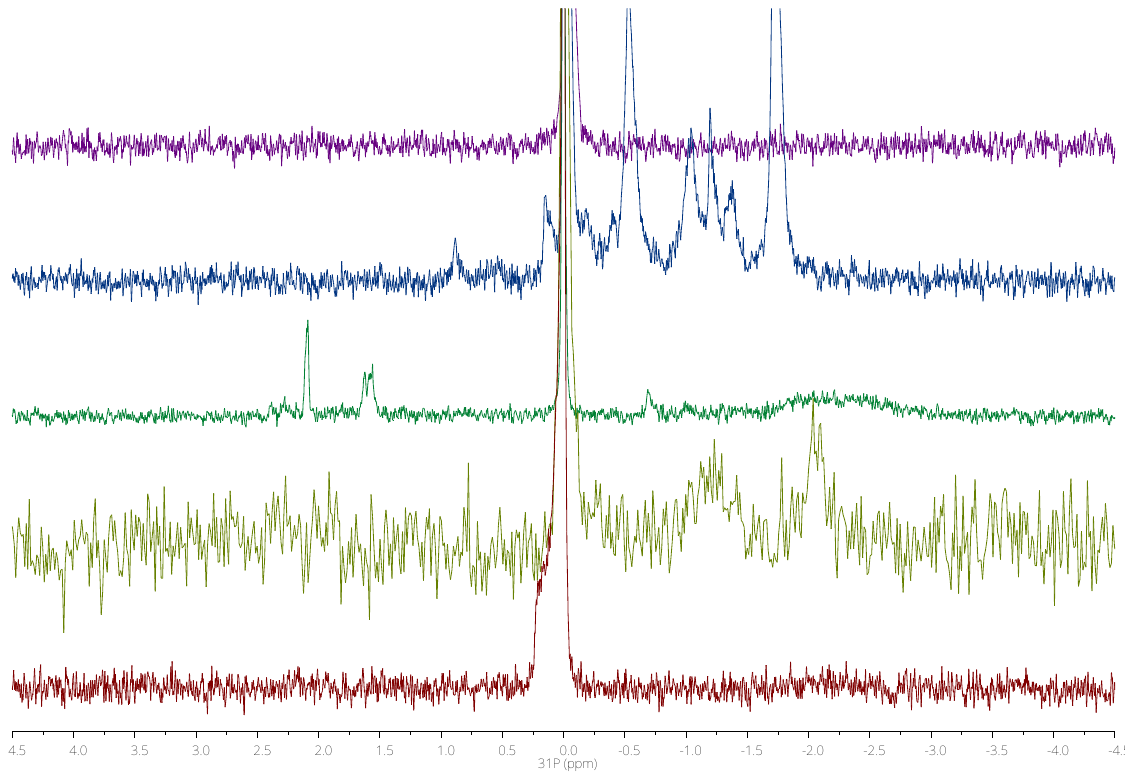

**Supplementary Figure 2:** (A) ^31^P NMR spectrum of H_3_PO_4_ solution in EMIM-Ac ( **─** upper), asolectin standard in EMIM-Ac ( **─** second upper), SDS and lysozyme-treated *Pseudomonas aeruginosa* wild-type PAO1 planktonic cells ( **─** middle), lyophilized *Pseudomonas aeruginosa* wild-type PAO1 biofilm in EMIM-Ac (10 mg/mL) ( **─** second lower) and PAO1 planktonic cells in 1x PBS ( **─** lower) at 25°C, showing phospholipid peaks for asolectin in EMIM-Ac, for lysed *P. aeruginosa* cells (SDS, lysozyme) in water, and the absence of phospholipid peaks in the spectrum of *P. aeruginosa* treated with EMIM-Ac. (B) Recovery yield of calf thymus DNA, lipase and cytochrome c standards following EMIM-Ac solubilization and recovery with perchloric acid (upper) followed by ethanol (lower). Error bars indicate standard deviation.

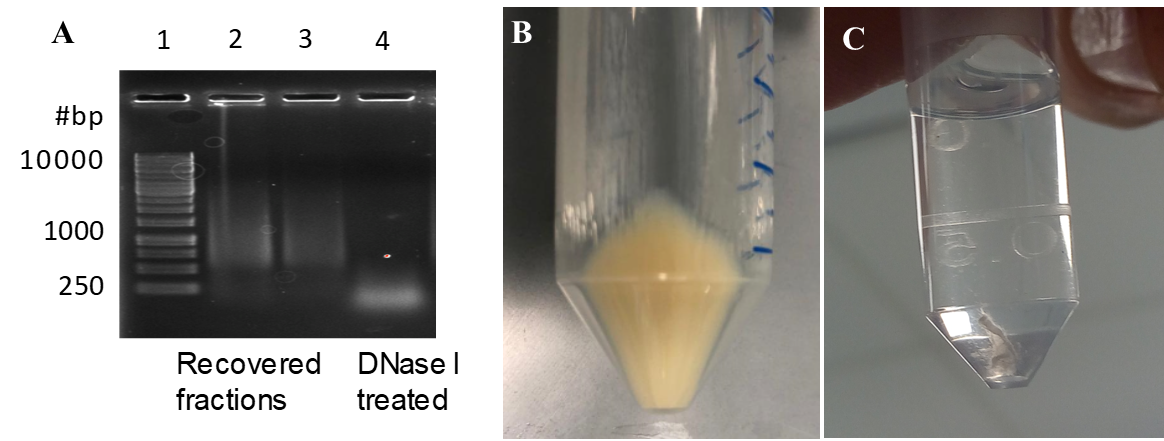

**Supplementary Figure 3:** (A) Agarose electrophoretic gel showing *P. aeruginosa* eDNA (Lanes 2 and 3) and digested with DNase (Lane 4). Lane 1 is the GeneRuler 1 kbp ladder. (B) Photograph of *P. aeruginosa* biofilm extracellular polymeric substances recovered by ethanol precipitation (70 % v/v) from 1-ethyl-3-methylimidazolium, dialyzed against double distilled water and then pelleted by centrifugation (10,000 g, 4°C) after it was dissolved (20 mg.mL^-1^) and eDNA first removed by perchloric acid precipitation (5 % v/v), showing that gelation was not observed once the nucleic acids were removed. (C) Photograph of calf thymus DNA recovered from 1-ethyl-3-methylimidazolium after dissolution (6 mg.mL^-1^), perchloric acid precipitation and dialysis against double distilled water showing that gelation is not a universal feature of all DNA following dissolution in 1-ethyl-3-methylimidazolium acetate and perchloric acid precipitation.

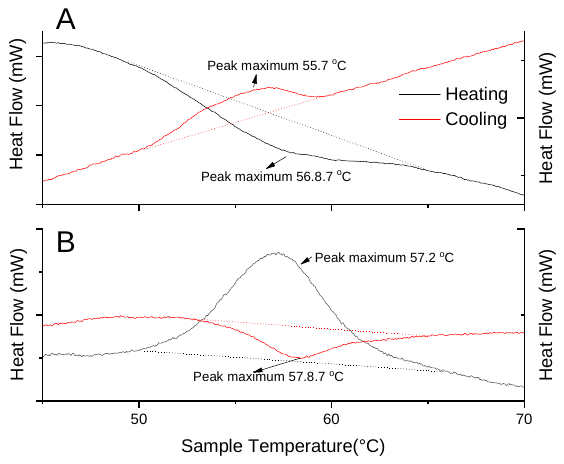

**Supplementary Figure 4: Differential scanning calorimetry (DSC) thermograms of (A) *Pseudomonas aeruginosa* wild type biofilm and (B) eDNA gel isolate showing phase transition peaks in heating (black) and cooling (red) .**

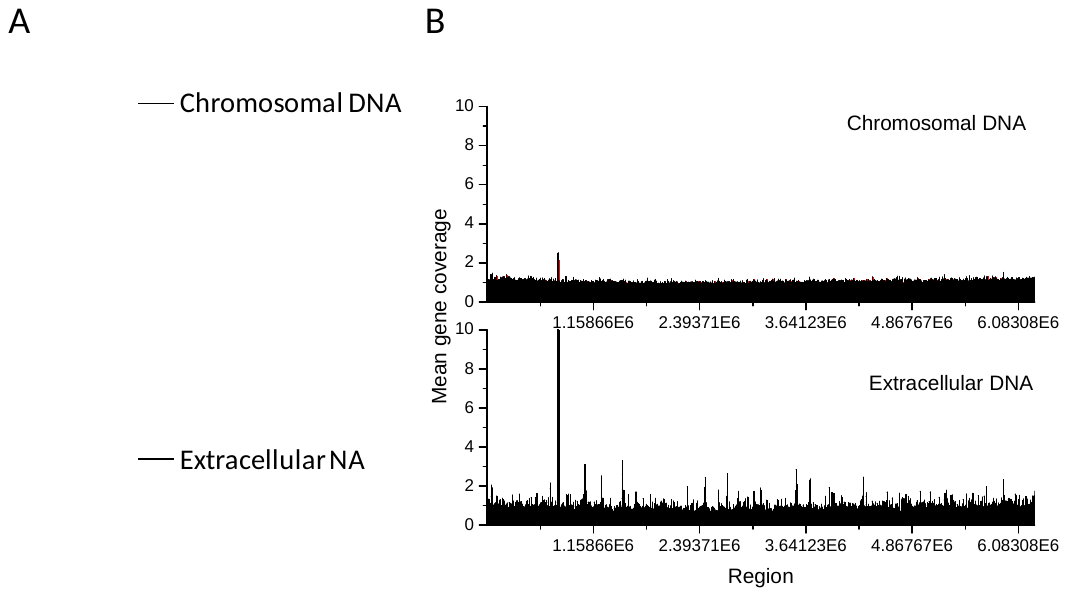

**Supplementary Figure 5:** Agarose gel loaded with chromosomal DNA extracted from *P. aeruginosa* pre-culture planktonic cells (upper) and extracellular NA gel isolate (lower) (A). Gene coverage of *P. aeruginosa* biofilm chromosomal (upper) and extracellular (lower) DNA normalized against *rpoB* numbers. The red oval denotes the peak resulting from bacteriophage Pf1 genes (B).

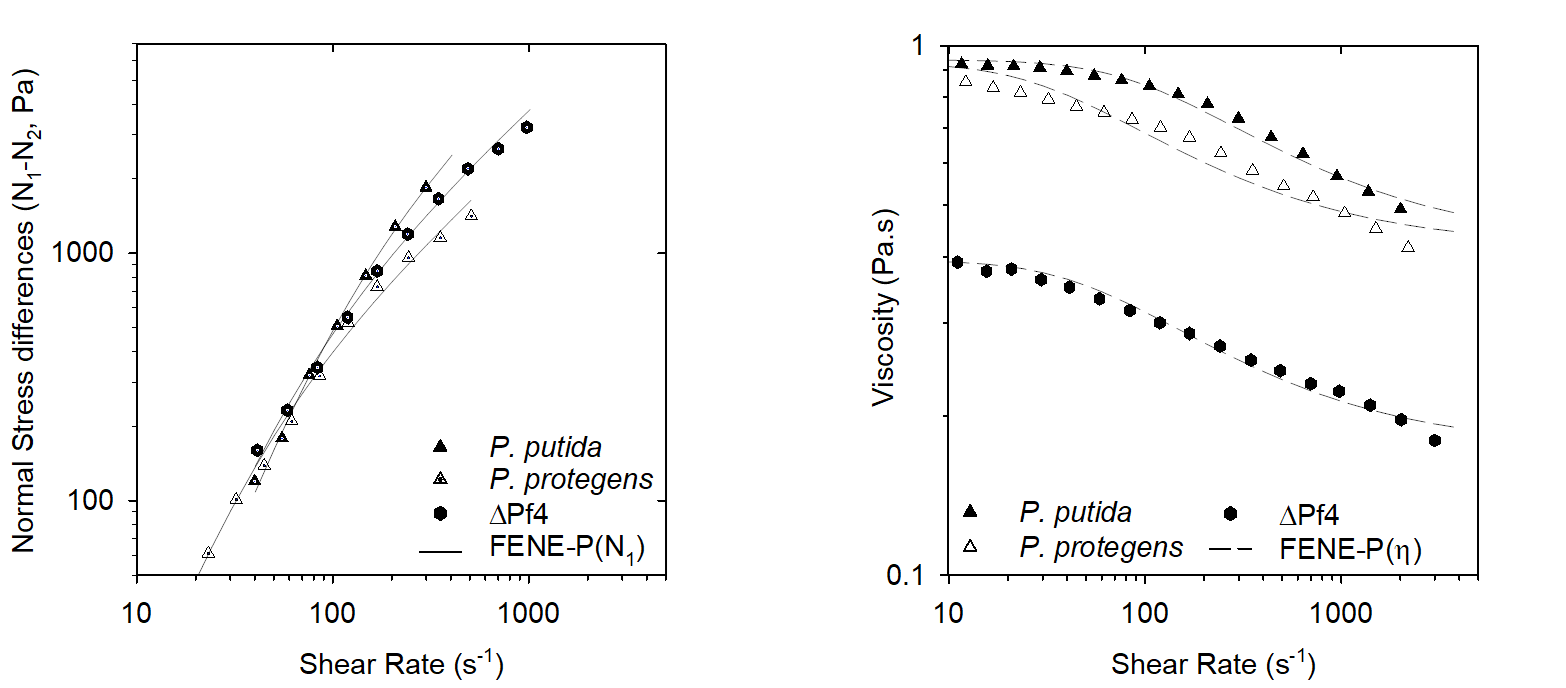

**B**

**A**

**Supplementary Figure 6:** (A) Normal stress differences (*N_1_ –N_2_*) and (B) shear stress as a function of shear rate for *Pseudomonas* biofilms: *P. putida, P. protogens* and *P. aeruginosa* ΔPf4, dissolved in 1-ethyl-3-methylimidazolium acetate (40 mg.mL^-1^) at 25 °C. Lines indicate FENE-P fits to the data.

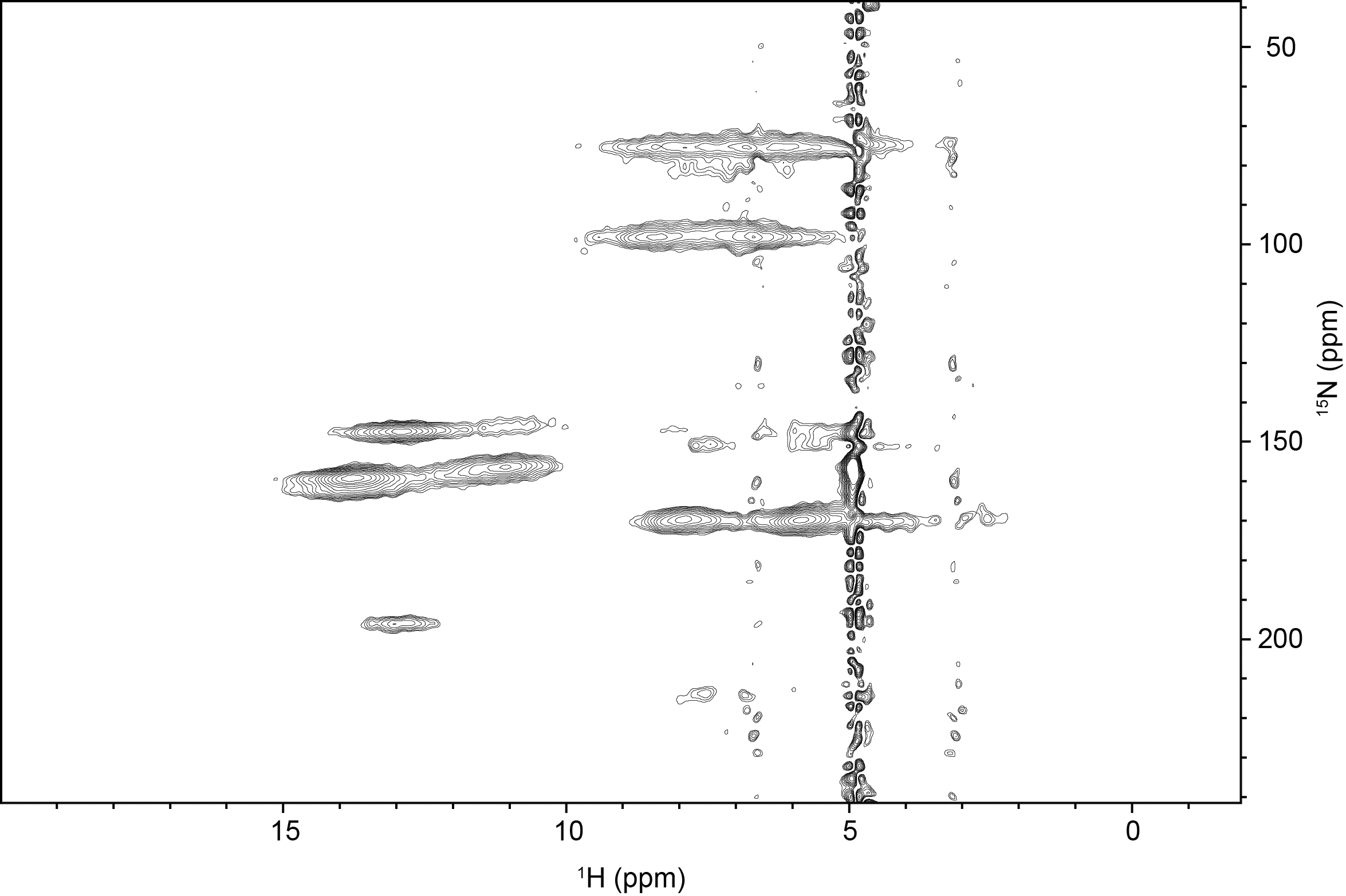

**Supplementary Figure 7:** Representative solid-state 2D ^1^H-^15^N through-space heteronuclear correlation (HETCOR) spectrum of extracellular nucleic acid (NA) gel isolate in double distilled water (2 mg), T = 25°C.

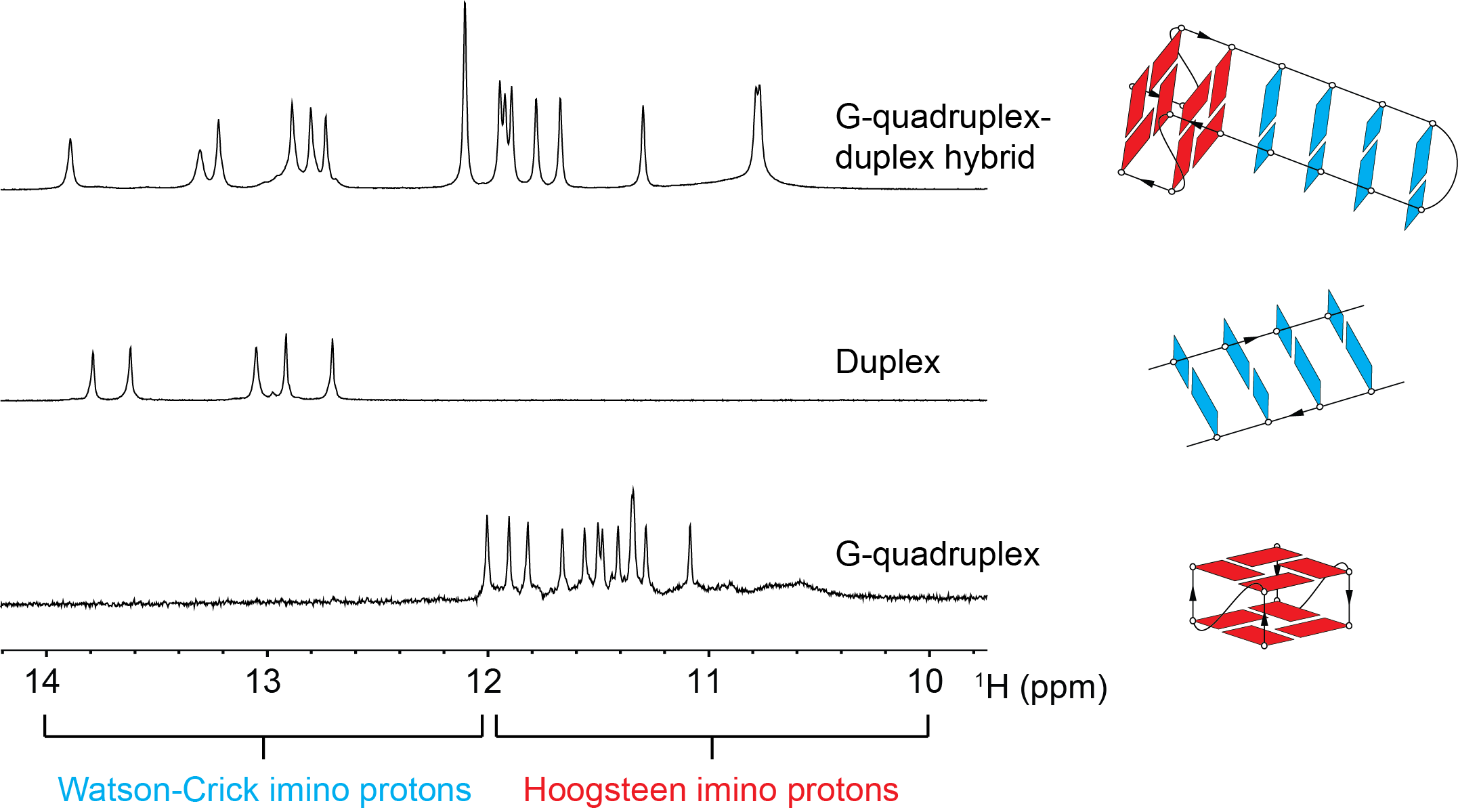

**B**

**C**

**A**

**Supplementary Figure 8:** 1-D ^1^H NMR spectra in the imino proton region of well-characterized (A) G-quadruplex (1), duplex (2) and quadruplex-duplex hybrid (3) structures. The 10-12 ppm region shows Hoogsteen-bonded imino protons from non-canonical base-pairs, while the 12-14 ppm region shows Watson-Crick-bonded imino protons from canonical base-pairs. The schematics on the right are for illustration purposes only.

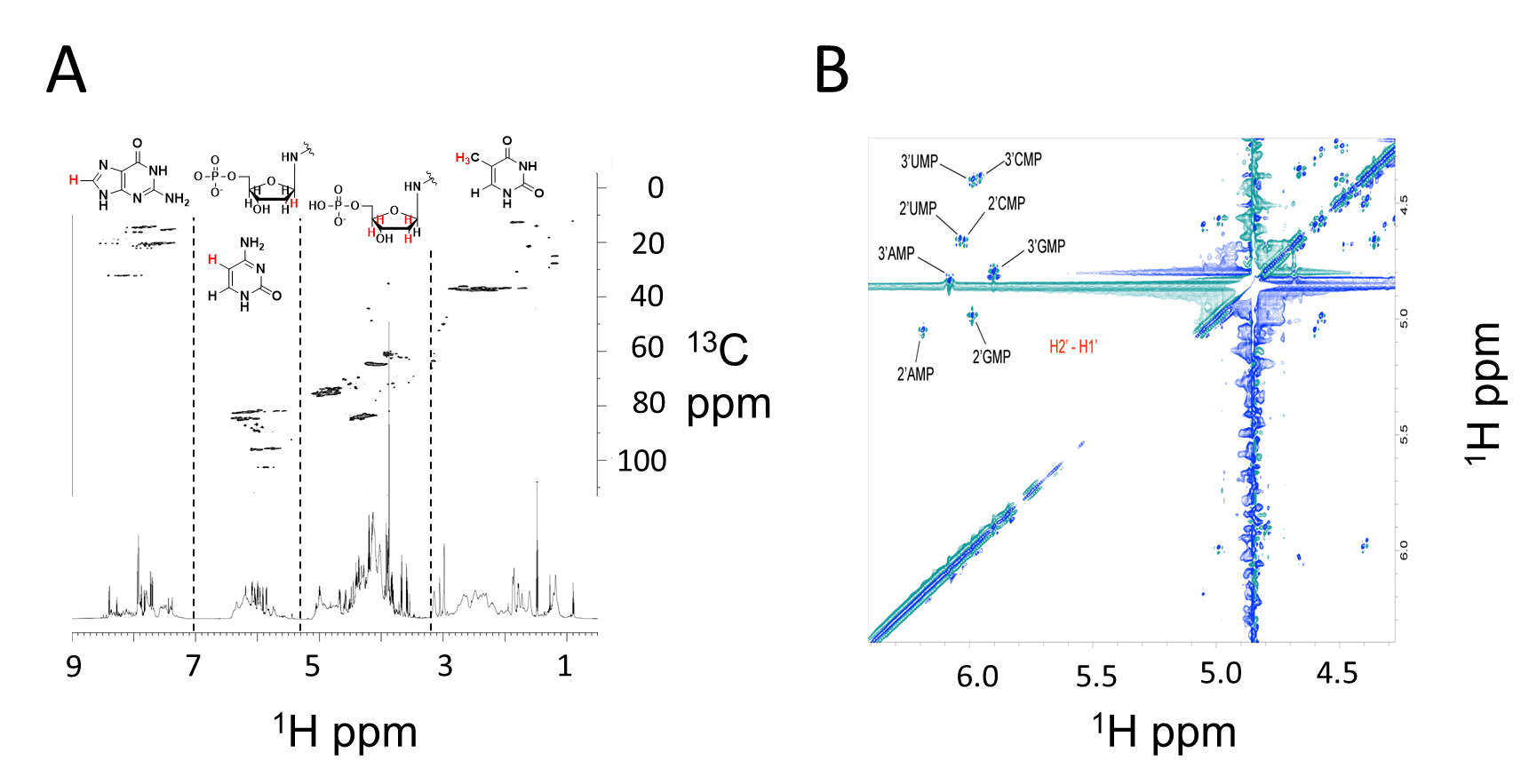

**Supplementary Figure 9 (**A) ^1^H-^13^C HSQC and 1-D ^1^H NMR spectra of *P. aeruginosa* biofilm extracellular NA gel isolate following alkalinization (0.1 M NaOD, 10 mg.mL^-1^, 55°C, 2 h) at 25°C showing a distribution of ^1^H-^13^C HSQC-TOCSY cross peaks that is consistent with the presence of nucleic acids and the absence of proteins and hexose-based sugars. (B) ^1^H-^1^H COSY NMR spectrum of *P. aeruginosa* biofilm extracellular NA gel isolate following alkalinization (0.1 M NaOD, 10 mg.mL^-1^, 55°C, 2 h).

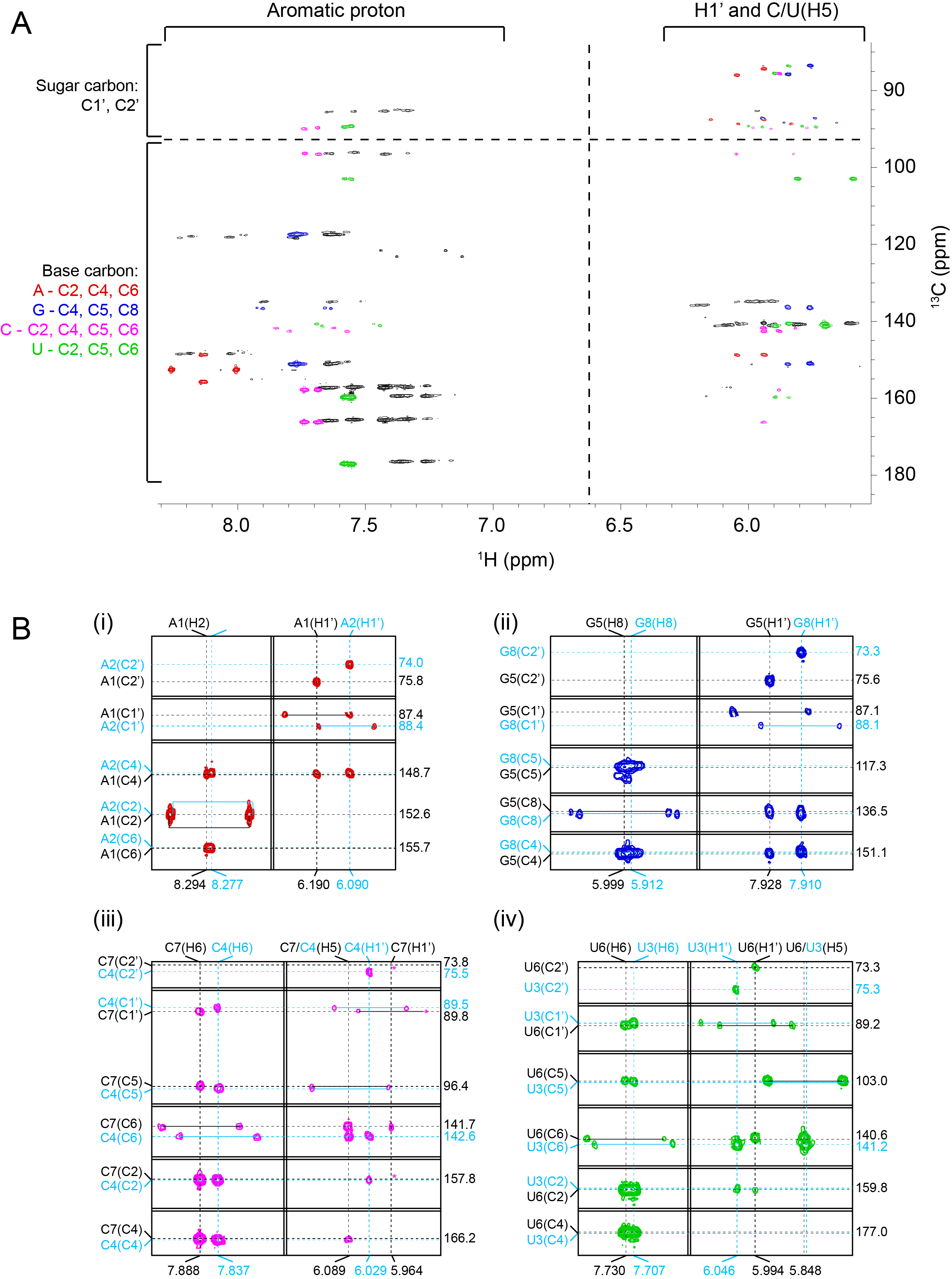

**Supplementary Figure 10:** (A) ^1^H-^13^C Heteronuclear multiple bond correlation (HMBC) NMR spectrum of *P. aeruginosa* biofilm extracellular NA gel isolate at 25°C following alkalinization (0.1 M NaOD, 10 mg.mL^-1^, 55°C, 2 h). The spectrum is divided to four regions with dashed lines corresponding to the observed interactions between sugar carbons to sugar protons, sugar carbons to base protons, base carbons to sugar protons and base carbons to base protons. Cross-peaks from monoribonucleotides are uniquely color coded. (B) The detailed assignments based on standard chemical shift values of ribonucleic acids, (i) adenine, (ii) guanine, (iii) cytosine and (iv) uracyl. All the assigned cross peaks are marked; the pairs of cross peaks connected by solid lines represents coupled one-bond carbon-proton interactions.

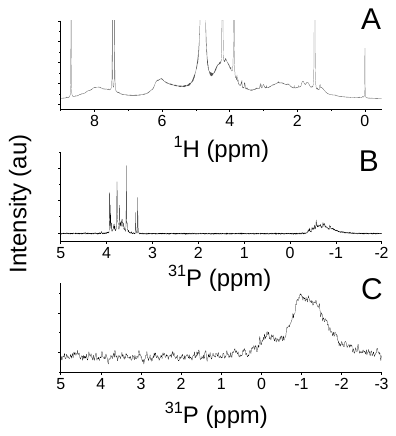

**Supplementary Figure 11:** (A)^1^H NMR spectra of *P. aeruginosa* biofilm NA gel isolate in D_2_O at 25°C. (B) ^31^P NMR spectrum of RNA standard (torula yeast) at 25°C following alkalinization (0.1 M NaOD, 10 mg.mL^-1^, 55°C, 2 h) and (C) in D_2_O following EMIM-Ac solubilisation and perchloric acid recovery (10 mg.mL^-1^) demonstrating that alkalinization only leads to RNA transesterification.

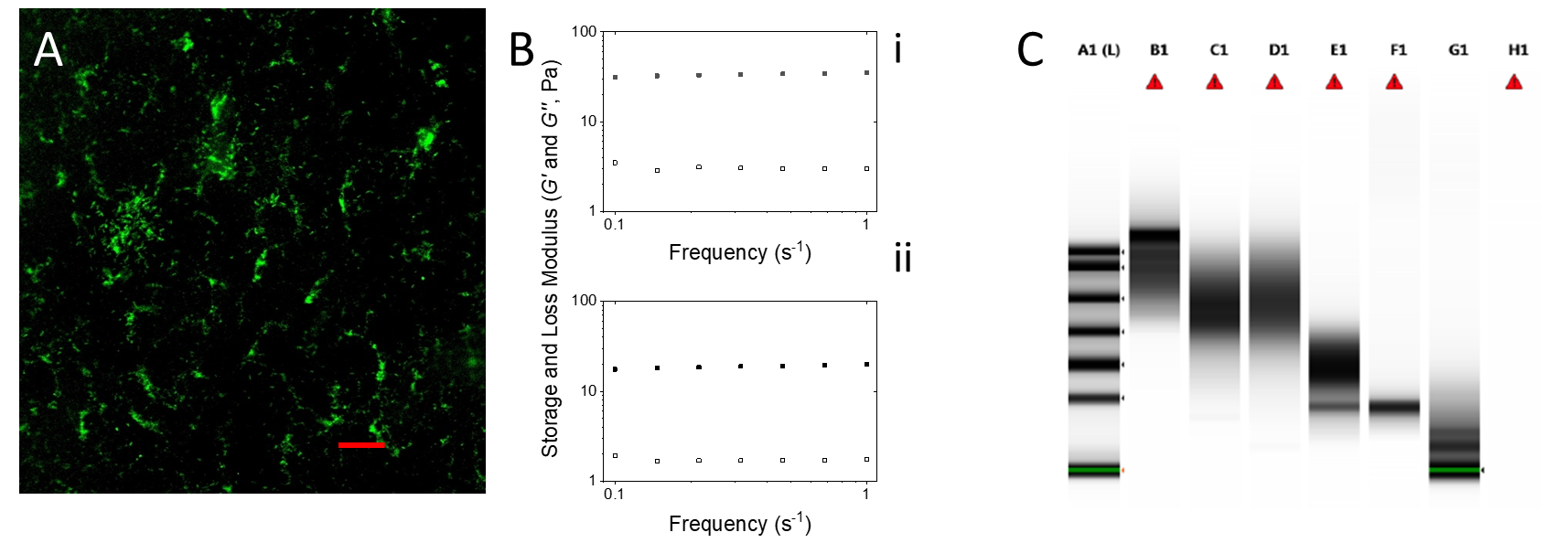

**Supplementary Figure 12:** (A) Micrograph of *P. aeruginosa* NA gel isolate, digested with DNase I and stained green with SYTO RNASelect, showing that SYTO RNASelect is binding to extracellular RNA (scale bar 10 µm). (B) Storage modulus (*G’*) and loss modulus (*G’’*) of *P. aeruginosa* wild type biofilm extracellular nucleic acid gel isolate in frequency sweep at 25°C, 250 µm gap, 0.1% strain, showing that the biofilm behaves like a gel at low amplitude (i.e. *G’* > *G’’*) even after digestion with RNase III (i) and RNase H (ii). (C) RNA ScreenTape® analysis of *Pseudomonas aeruginosa* wild type eDNA isolate without pre-heating (Lane B1), after pre-heating to 60 °C (Lane C1), digestion with RNase A (Lane D1), digestion with DNase I (Lane E1), digestion with DNase I followed by RNase A (Lane F1). A1 is nucleobase ladder, from top 6000, 4000, 2000, 1000, 500, 200 and 25 nucleotides.

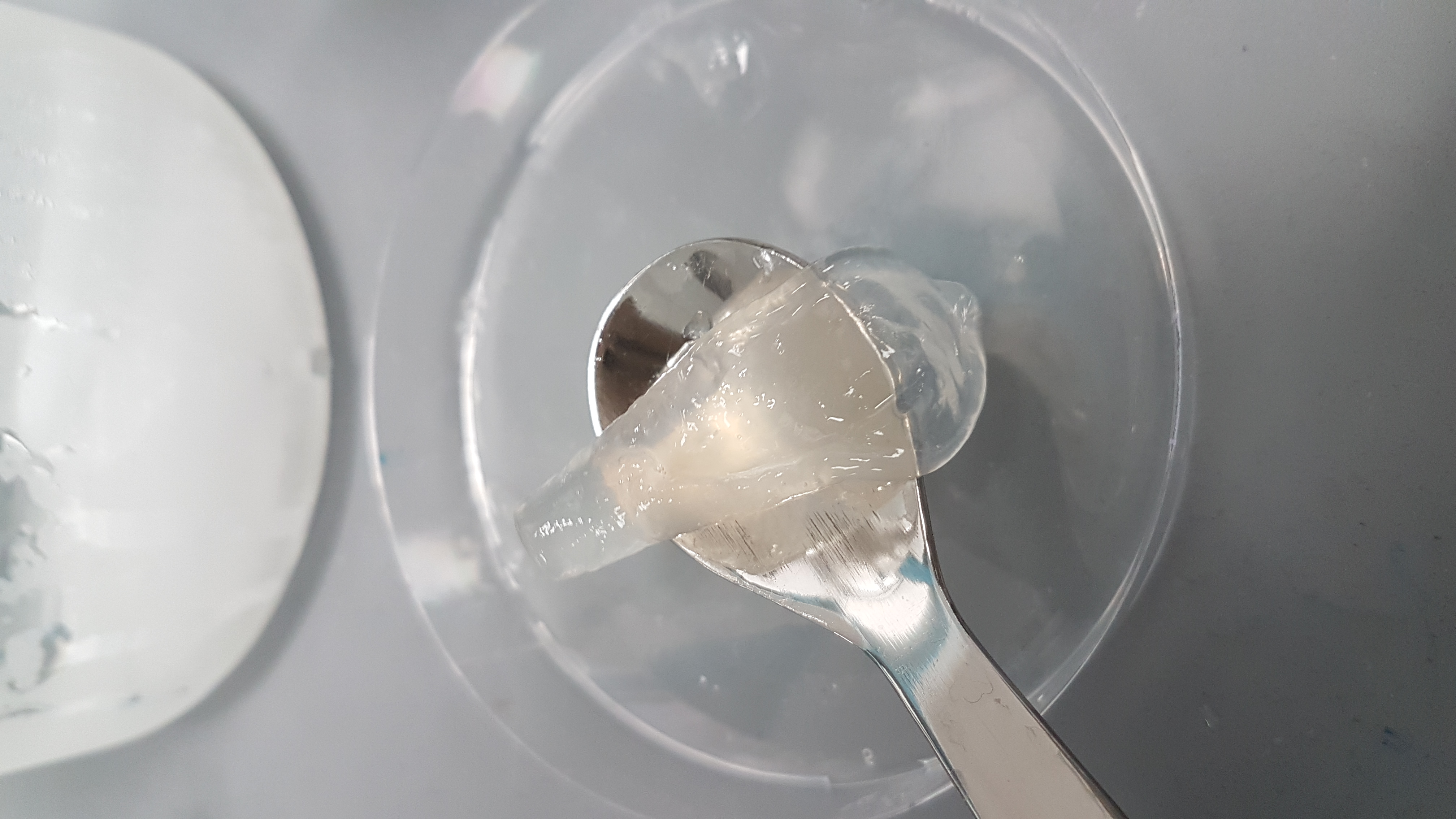

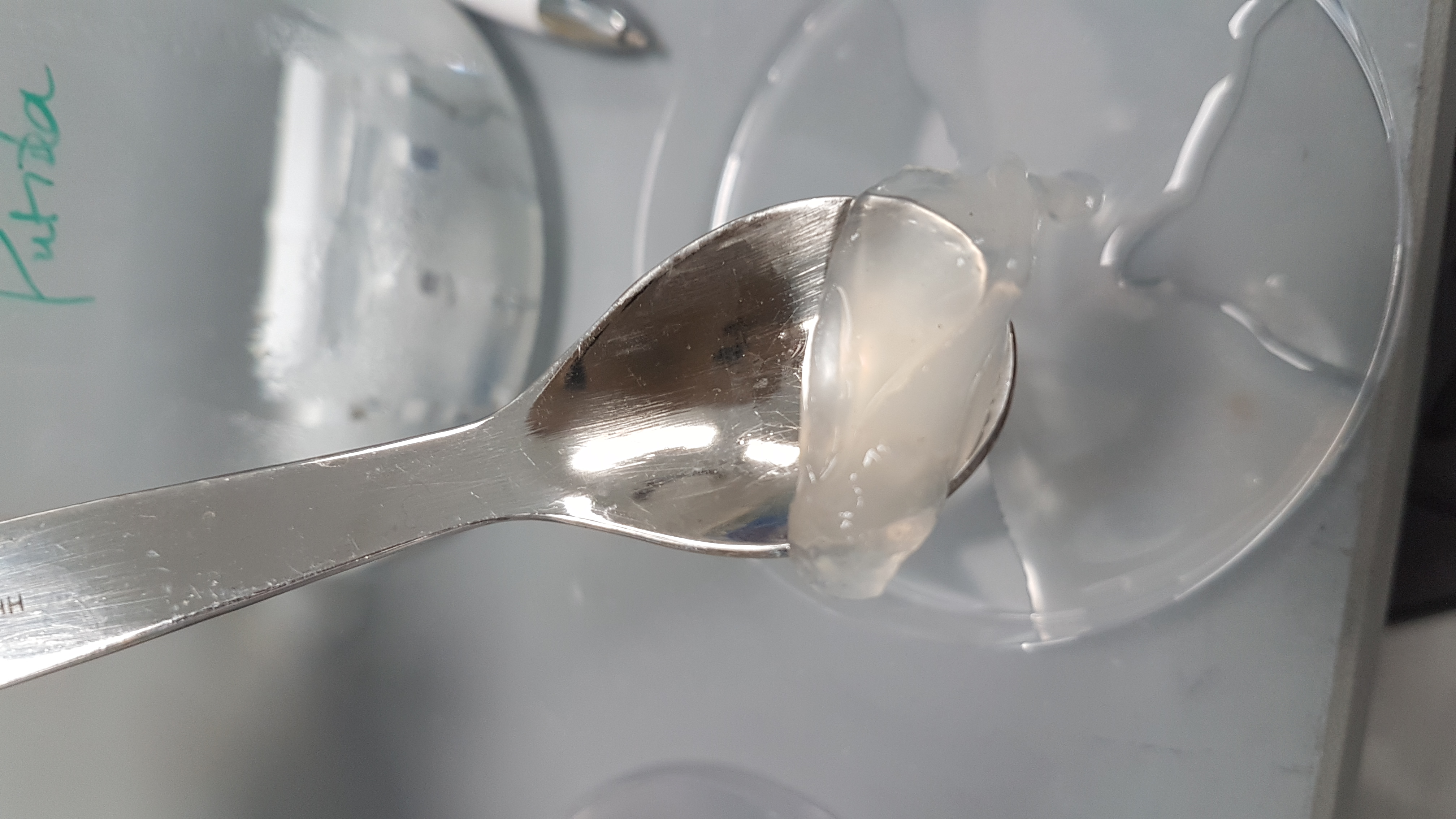

**B**

**A**

**Supplementary Figure 13:** Photograph of nucleic acid gel extracted from (A) *P. protegens* and (B) *P. putida*.

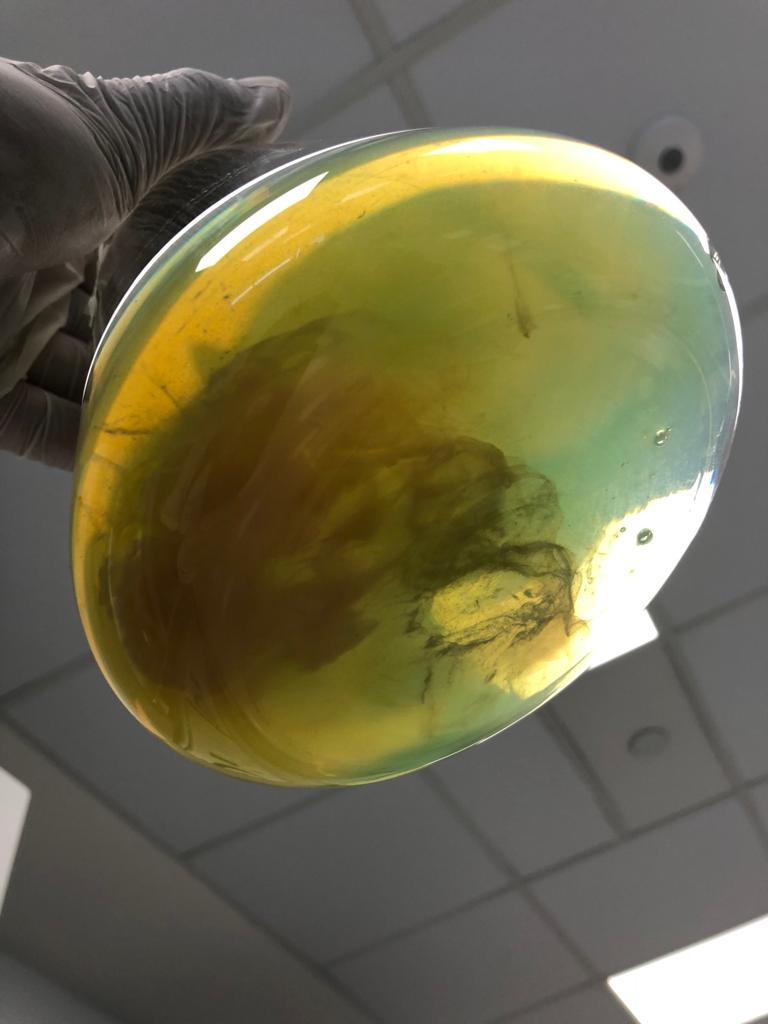

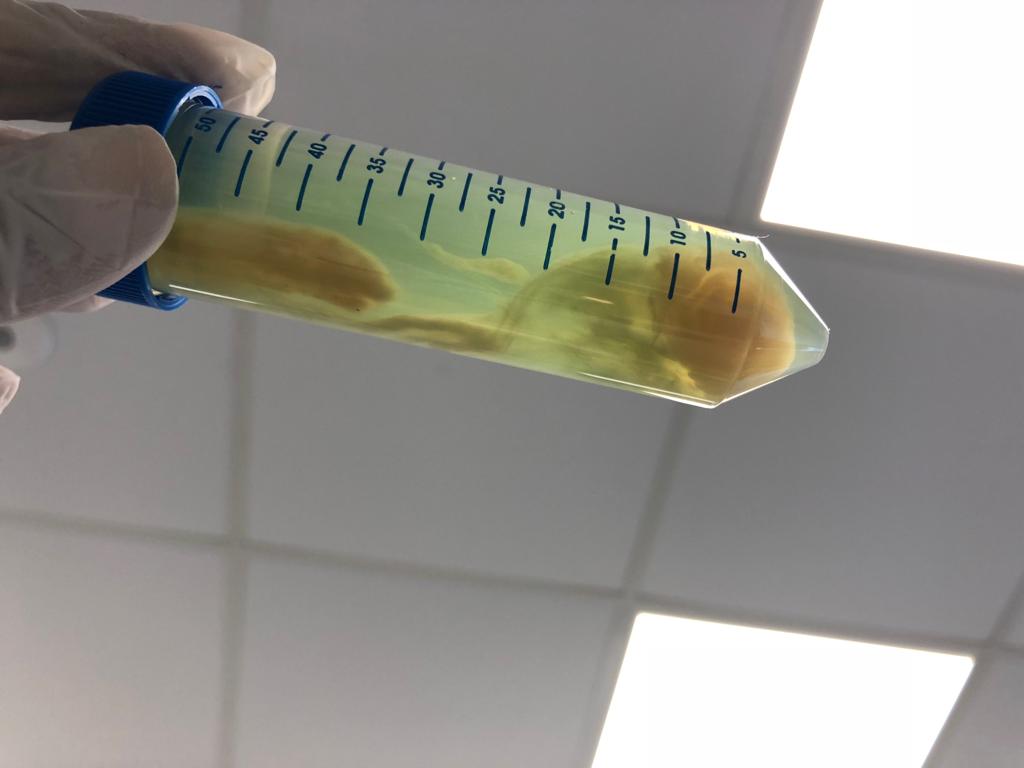

**B**

**A**

Supernatant

Biofilm

**Supplementary Figure 14:** Photographs of *P. aeruginosa* biofilm after 5-d static growth in lysogeny broth medium at 37°C in 2 L Erlenmeyer flask (A) before centrifugation and (B) in 50 mL centrifuge tube after centrifugation showing the separation of biofilm from supernatant.

**Supplementary Table 1:** Power law exponent (i.e. *n* in $N_{1}=K\dot{\gamma}^{n}$ and *m* in $\sigma=K_{\sigma}\dot{\gamma}^{m}$) of *P. aeruginosa* biofilms following dissolution in EMIM-Ac. *m* values approaching unity indicate a Newtonian-like fluid property. Viscosity is slightly shear thinning (*m* = 0.8 to 0.9) for all samples except RNase, DNase and EMIM-Ac which are Newtonian like (*m* ≥ 0.93), which would be expected from dilute polymer solutions in viscous fluids (i.e. Boger fluids).

|  | $\sigma=K_{\sigma}\dot{\gamma}^{m}$ | | | | $N_{1}=K_{N1}\dot{\gamma}^{n}$ | | | |
| --- | --- | --- | --- | --- | --- | --- | --- | --- |
| **Sample** | **K_σ_** | | ***m*** | | **K_N1_** | | ***n*** | |
|  | **Ave.** | **Std Dev** | **Ave.** | **Std Dev** | **Ave.** | **Std Dev** | **Ave.** | **Std Dev** |
| *P. aeruginosa* wild type | 0.63 | 0.08 | 0.84 | 0.01 | 1.44 | 0.45 | 1.36 | 0 |
| ΔPsl | 1.26 | 0.35 | 0.76 | 0.04 | 7.81 | 5.18 | 1.21 | 0.15 |
| ΔPel | 0.35 | 0 | 0.89 | 0 | 0.58 | 0.15 | 1.34 | 0.02 |
| Pronase | 0.55 | 0.03 | 0.89 | 0 | 0.18 | 0.11 | 1.64 | 0.08 |
| RNase | 0.34 | 0.01 | 0.93 | 0 | 2.11 | 0.09 | 0.91 | 0.01 |
| *P. putida* | 1.32 | 0.01 | 0.91 | 0.01 | 0.56 | 0.16 | 1.48 | 0.08 |
| PDO300 | 0.73 | 0.35 | 0.89 | 0.01 | 2.33 | 0.59 | 1.10 | 0.03 |
| *P. protegens* | 1.14 | 0.01 | 0.90 | 0.01 | 2.75 | 1.30 | 1.10 | 0.11 |
| ΔPf4 | 0.57 | 0.02 | 0.87 | 0.01 | 1.59 | 1.25 | 1.33 | 0.24 |
| DNase | 0.20 | 0.01 | 0.99 | 0.01 | - | | - | |
| EMIM-Ac | 0.13 | 0 | 0.98 | 0 | - | | - | |

**Supplementary Table 2:** Fitting parameters for the FENE-P model including λ_1_ = relaxation time, b = a measure of the relative extensibility of the model spring, η_s_ = solvent viscosity, η_p_ = polymer contribution to the viscosity. Molecular extensibility and relaxation times, as predicted by FENE-P, decrease in accordance with elasticity.

|  | **Average** | | | |  | **Std. Deviation** | | | |
| --- | --- | --- | --- | --- | --- | --- | --- | --- | --- |
| **Sample** | b | λ_1_ | η_p_ | η_s_ |  | b | λ_1_ | η_p_ | η_s_ |
| *P. aeruginosa* wild type | 2248.5 | 0.322 | 0.260 | 0.145 |  | 643.1 | 0.066 | 0.040 | 0.0026 |
| ΔPsl | 3511.5 | 0.676 | 0.381 | 0.161 |  | 1000.9 | 0.219 | 0.113 | 0.0114 |
| ΔPel | 2000.8 | 0.227 | 0.110 | 0.133 |  | 637.8 | 0.056 | 0.000 | 0.0025 |
| Pronase | 1871.8 | 0.103 | 0.197 | 0.167 |  | 693.1 | 0.047 | 0.015 | 0.0339 |
| RNase | 399.2 | 0.102 | 0.095 | 0.183 |  | 17.6 | 0.008 | 0.003 | 0.0077 |
| PDO300 | 909.4 | 0.453 | 0.138 | 0.147 |  | 134.2 | 0.023 | 0.026 | 0.0007 |
| DNase | 135.7 | 0.005 | 0.020 | 0.129 |  | 106.8 | 0.001 | 0.011 | 0.0112 |
| ΔPf4 | 865.4 | 0.230 | 0.243 | 0.166 |  | 129.9 | 0.034 | 0.022 | 0.0072 |
| *P. protegens* | 137.2 | 0.175 | 0.501 | 0.414 |  | 19.2 | 0.028 | 0.025 | 0.0013 |
| *P. putida* | 336.9 | 0.071 | 0.544 | 0.440 |  | 53.5 | 0.002 | 0.018 | 0.0199 |
